# Pulmonary stromal expansion and intra-alveolar coagulation are primary causes of Covid-19 death

**DOI:** 10.1101/2020.12.23.424172

**Authors:** Laszlo Szekely, Bela Bozoky, Matyas Bendek, Masih Ostad, Pablo Lavignasse, Lars Haag, Jieyu Wu, Xu Jing, Soham Gupta, Elisa Saccon, Anders Sönnerborg, Yihai Cao, Mikael Björnstedt, Attila Szakos

## Abstract

Most Covid-19 victims are old and die from unrelated causes. Here we present **t**welve complete autopsies, including two rapid autopsies of young patients where the cause of death was Covid-19 ARDS. The main virus induced pathology was in the lung parenchyma and not in the airways. Most coagulation events occurred in the intra-alveolar and not in the intra-vascular space and the few thrombi were mainly composed of aggregated thrombocytes. The dominant inflammatory response was the massive accumulation of CD163+ macrophages and the disappearance of T killer, NK and B-cells. The virus was replicating in the pneumocytes and macrophages but not in bronchial epithelium, endothel, pericytes or stromal cells. The lung consolidations were produced by a massive regenerative response, stromal and epithelial proliferation and neovascularization. We suggest that thrombocyte aggregation inhibition, angiogenesis inhibition and general proliferation inhibition may have a roll in the treatment of advanced Covid-19 ARDS.

## Introduction

Ten months in the Covid-19 pandemic with close to 50 million of confirmed cases and over one million deaths worldwide the exact mechanism of the Sars-Cov2 contribution to death of Covid-19 victims is still only partially understood. The major impediment is the lack of high-quality rapid autopsies from the informative cases that can be used to discern the exact role of the virus in the induced pathology. The available reports with autopsy data are either based on autopsy findings of small number of cases or summarize findings of unselected elderly population with numerous co-morbidities (Zhou et al., 2020)(Calabrese et al., 2020)(Bradley et al., 2020)(Edler et al., 2020)(Wichmann et al., 2020)(Barton et al., 2020). As the median age of people who died while infected with Sars-Cov2 is close to eighty-year-old and most of them had a few serious co-morbidities, the exact role of the virus in the complex process of health deterioration is difficult to discern.

Sweden was hit hard at the beginning of the pandemic with 85000 confirmed cases and 5800 deaths by 2020 end of August peaking in April. Close to 90% of the victims were aged over 70 and almost 50% over 85. Only one percent was under the age of 50 and only 3.7 percent under the age of 60. Many victims also received ICU care with extensive list of medical interventions including prolonged overpressure respiration. Virus induced pathology is a result of a complex combination of events, including virus induced direct cytopathic effects on the infected cells, bystander effect on non-infected cells, systemic non-cellular effect of the coagulation and complement system, regenerative and immune response of the host and additional pathologies exerted by other opportunistic pathogens that take advantage of damaged barrier structures or weakened innate or adoptive defenses.

To effectively dissect the pathogenic role of the virus requires detailed examination of selected cases with minimal co-morbidities and limited post-mortem artefacts. Although the Koch’s time for Covid-19, the time required from identification of a new infectious disease until the submission of the genome sequence of its pathogen to public database was unprecedently short, 15 days (Chen et al., 2020) – the way how and why the virus causes disease and occasionally kills the host is much more difficult to elucidate. High quality autopsies with detailed histological and immunohistochemical follow-up are therefore essential (Salerno et al., 2020)(Kowalik et al., 2020)l.

The available autopsy data and experimental results on ex vivo infected material suggested droplet-based transmission that primarily infects upper airway epithelial cells, rich in ACE2 viral receptor and subsequent downward spreading to the lower airways in the severe cases (Hui et al., 2020). RT-PCR based detection of the viral RNA from multiple organs raised the possibility of extensive additional virus induced organ damage (Hanley et al., 2020). The main pulmonary pathology has been defined as diffuse alveolar damage (DAD) and subsequent acute respiratory distress syndrome (ARDS) with severe and frequent thromboembolic complications with intravascular fibrin deposition, for review see (Vasquez-Bonilla et al., 2020). The emphasis on the presence of fibrin micro-thrombi prompted vigorous anti-coagulative therapies. Recent meta-analysis of the published clinical data however showed no significant increased survival of patients who received anti-coagulative therapy (Salah et al., 2020)

Here we present a series of detailed autopsy findings of relatively young individuals where the direct cause of death was Covid-19 induced ARDS. We carried out extensive histological, immunohistochemical and virus RNA in situ analysis on single cell levels on cases with short postmortem elapsed time. Our findings are only partially congruent with previous reports and suggest alternative therapeutic options for severe cases.

## Results

### Patient population

In this report we include data of 12 patients where the direct cause of death was determined as COVID-19 associated ARDS. The patient characteristics are summarized in Table I. Briefly the median age of the group was 58 years (from 23 to 81) and the median BMI 26.8. The median time from the first appearance of the symptoms until death was 13 days. Five patients were treated at the ICU unit and received mechanical ventilation. Eight patients were subjected to CPR procedure immediately prior death. Three of them were treated with the automatic chest compression device of LUCAS. Four patients died at home. Seven patients had high blood pressure in the anamnesis, five of them also suffered from Type II diabetes. One patient was suffering from morbid obesity. Four patients had no previously identified risk conditions.

Patients characteristic are in Table 1. Organ weights are summarized in Table 2. All patients showed greatly increased serum CRP. Most showed increased levels of D-dimer (7/8) and troponin (7/10) as well.

### Gross autopsy and histology findings

#### External examination

The autopsies were carried out shortly after the death (median 6 days). Two autopsies were rapid autopsies (within 9 (C11) respectively 15 (C9) hours and two within 2 (C1) respectively 3 (C8) postmortem days. Even with the longest postmortem period (14 days) most organs showed well preserved macroscopic morphology.

External examination of the bodies showed age adequate cutaneous conditions and revealed no exanthemas, bleedings or other viral infection associated cutaneous changes. Occasional subcutaneous edema (C4, C9) pemphigoid (C5) and chronic venous stasis associated induration on the skin of the leg (C9) was noted.

#### Respiratory system

Upon internal examination the most prominent macroscopic pathological changes were observed in the lungs. The pleural surfaces were regularly smooth and glossy with patchy hemorrhagic or pale discolorations that sharply respected the lobular borders. There was modest amount of accumulation of serous or bloody pleural effusion fluids bilaterally with median volume of 85 ml (IQR 30-150 ml). There was no macroscopically obvious purulent or acute adhesive pleuritic inflammation in any of the cases. Occasional old fibrous adherences were detected. The lungs of all victims were heavy and fluid filled, approximately 2,8 times heavier than the corresponding average female and male lungs (Molina & DiMaio, 2015). All lungs were firm to touch with extensive rubbery consolidations. The consolidations were often dark colored indurations that respected the lobules borders (Figure 1). Whitish accentuation of the lobules borders was also often observed as with lung edema. All the lungs showed no or only minimal air content. The macroscopic changes correlated well with the CT images showing patchy, sharply demarcated ground glass homogenizations. The consolidated areas showed various levels of discoloration ranging from pale cream color to dark gray red, dependent on the amount of blood. The posterior part of the lower lobes was always free of air and fully consolidated, containing various amounts of blood, edema fluid and cellular elements. The lumens of most bronchi and blood vessels were regularly patent (Supplemental Figure 1)

**Figure 1.**
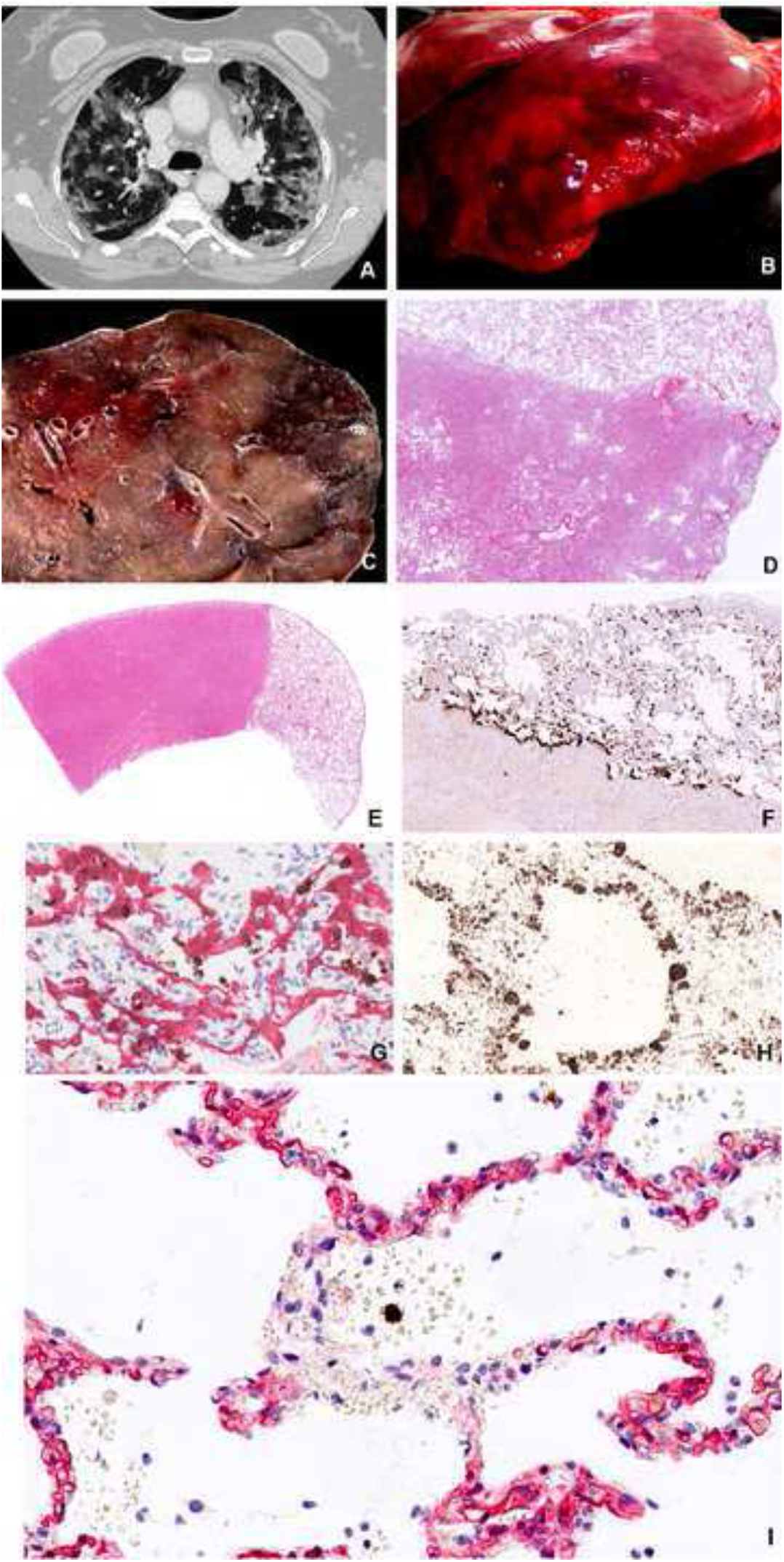
Characteristics of Covid-19 lungs. Consolidations in Covid-19 lungs show good correlation between the CT image (A), the macro image of the lung surface (B), the cut surface of the lower lobe (C) and the low magnification HE stained histology slide (D) showing fluid and cell filled areas that are sharply demarcated by the lobules border. (Patient C3). Sharply demarcated areas of mildly affected and massively consolidated lung areas shown by HE staining (E). Surfactant positive type II pneumocytes that are the primary site of viral replication are primarily associated with the mildly affected areas (F). Sars-Cov2 Spike RNA (brown signal) mainly associated with CK18 positive epithelial cells (red) as shown by combined RNAScope in situ hybridization and immunohistochemistry (G). The denuded alveolar walls are populated by CD68 positive macrophages in the consolidated area (H). Bystander damage of alveolar capillary wall (CD34 – red) in the vicinity of Sars-Cov2 infected alveolar cell (RNAScope Spike in situ - brown) that leads to intraalveolar bleeding. No infection of endothelial cells or pericyte is observable (I).

The mucous membranes of the upper and lower airways were regularly unaffected. Only two patients (C2 and C10) showed hemorrhagic mucous membranes in the upper airways (Supplemental Figure 2). Sample from C2 was free from inflammatory infiltrate whereas C10 suffered from purulent trachea-bronchitis. The sphenoid sinus was opened in every case to assess the inflammatory involvement of the intracranial mucous membranes. Only one patient (C2) showed blood filled swollen mucous membrane. The soft palate, uvula, tongue, pharynx, larynx, vocal cords, main and small bronchi were inspected, photographed and histologically sampled in order to attempt to demonstrate virus induced tissue damage. No macroscopically evident pathology was detected at any of these sites.

The lung damage was characterized by detailed immunohistochemical analysis. The most severe pathological changes occurred in the lung parenchyma starting with immense cytopathic effects on both Type I and Type II pneumocytes that were identified by Muc1 and surfactant immunostaining respectively. Upon infection the pneumocytes were greatly swollen and mobilized as single cells or confluent cell sheets. The damaged alveoli were regularly filled with edema fluid, plasma, coagulated fibrin, blood or coagulated blood. The mobilization of the pneumocytes was accompanied with cell proliferation as shown by Ki67 and CyclinD1 staining and concomitant expression of both type I and type II markers. The denuded alveolus wall was rapidly populated by CD68 positive macrophages (Figure 1).

The presence of virus replication was detected by RNA in situ hybridization using RNAScope Z probes against the Sars-Cov2 Spike RNA. Every Spike RNAScope slide was accompanied with a positive control slide with probe against ubiquitin mRNA and a negative control slide with a probe against the bacterial gene dapB. RNAScope was also combined with post hybridization immunostaining using cell type specific primary and alkaline phosphatase conjugated secondary antibodies to identify the virus bearing cells. The viral RNA most frequently localized to pneumocytes. Importantly bronchial epitheliums in the small and large bronchi, bronchial reserve cells, goblet cells were negative.

The desquamated pneumocytes often showed advanced cellular atypia with nuclear heterogeneity and bizarrely shaped cytoplasm. These atypical cells, colloquially called “Covid cells” were detectable in all the twelve lungs and along with the ubiquitous hyaline membranes, the histological markers of ARDS, constituted the most prominent histological hallmarks of the disease. Importantly only a small fraction of these cells were virus infected as shown by Spike RNA /cytokeratin 18 double labelling. Most bloated and desquamated epithelial cells showed various degree of vacuolar degeneration as shown by transmission electron microscopy without discernable production of mature virus particles (Figure 2).

**Figure 2.**
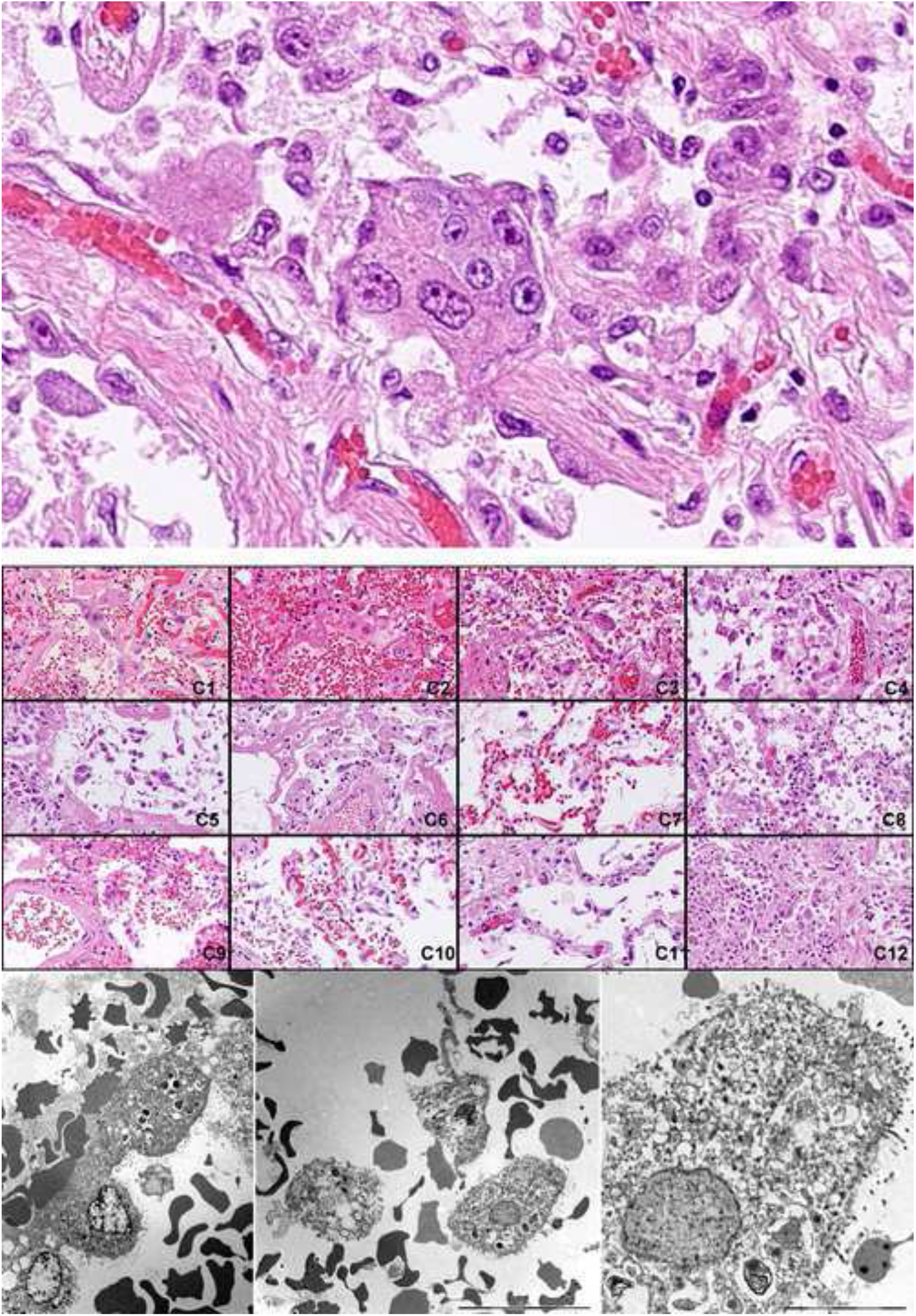
Identification of Covid cells. Desquamated atypical alveolar epithelial cells, “Covid cells” are one of the histological hallmarks of the virus induced lung damage.

Most viral replication has occurred in the morphologically mildly effected areas in cells that were CK18, CK19, EMA, surfactant A positives. Importantly up to a third of virus RNA signal was localized to cytokeratin negative, non-epithelial cells. Heavily consolidated areas contained manly disintegrated remnants of the infected cells with viral RNA positive debris. The denudation of the alveolar wall was regularly accompanied with the damage of the alveolar capillaries and release of plasma and eventually blood into the alveolar space. Importantly the endothelial cells did not show expression of viral RNA as shown by Spike RNA/CD34 double labelling. Spike RNA/CD68 double labelling revealed frequent presence of viral RNA in macrophages (Figure 6 J.) Similarly to the infected epithelial cells the viral RNA in situ signal in the CD68 positive cells ranged from distinct cytoplasmic granules to large confluent cytoplasmic fields filling up most of the cell volume. As unprotected RNA is rapidly degraded in phagosomes the presence of large confluent viral RNA represents actively replicating virus with amplified Spike subgenomic fragment. Even CD68 positive cell remnants with released virus RNA positive material were frequently encountered. Ingested non degradable particles in macrophages, as in anthracosis, might pose diagnostic difficulties to discriminate from true in situ signal. In our cases the RNAScope signal was readily distinguishable from the occasionally occurring anthracosis by its different color. Moreover, the anthracotic macrophages were present in the negative dapB control slides.

Infection of the alveolar epithelial cells is regularly accompanied with the damage of the alveolar wall capillaries in the vicinity of the infection event as illustrated in Figure 1 (I) and Figure 3 (A,B,C). The endothelial cells showed no sign of viral infection but various levels of cytopathic effects with formation of intracytoplasmic vacuoles and fenestration that led to plasma leakage as shown by the release of IgM and von Willebrand factor, intra-alveolar coagulation of the released fibrin or even frank bleeding with or without coagulation (Figure 3).

**Figure 3.**
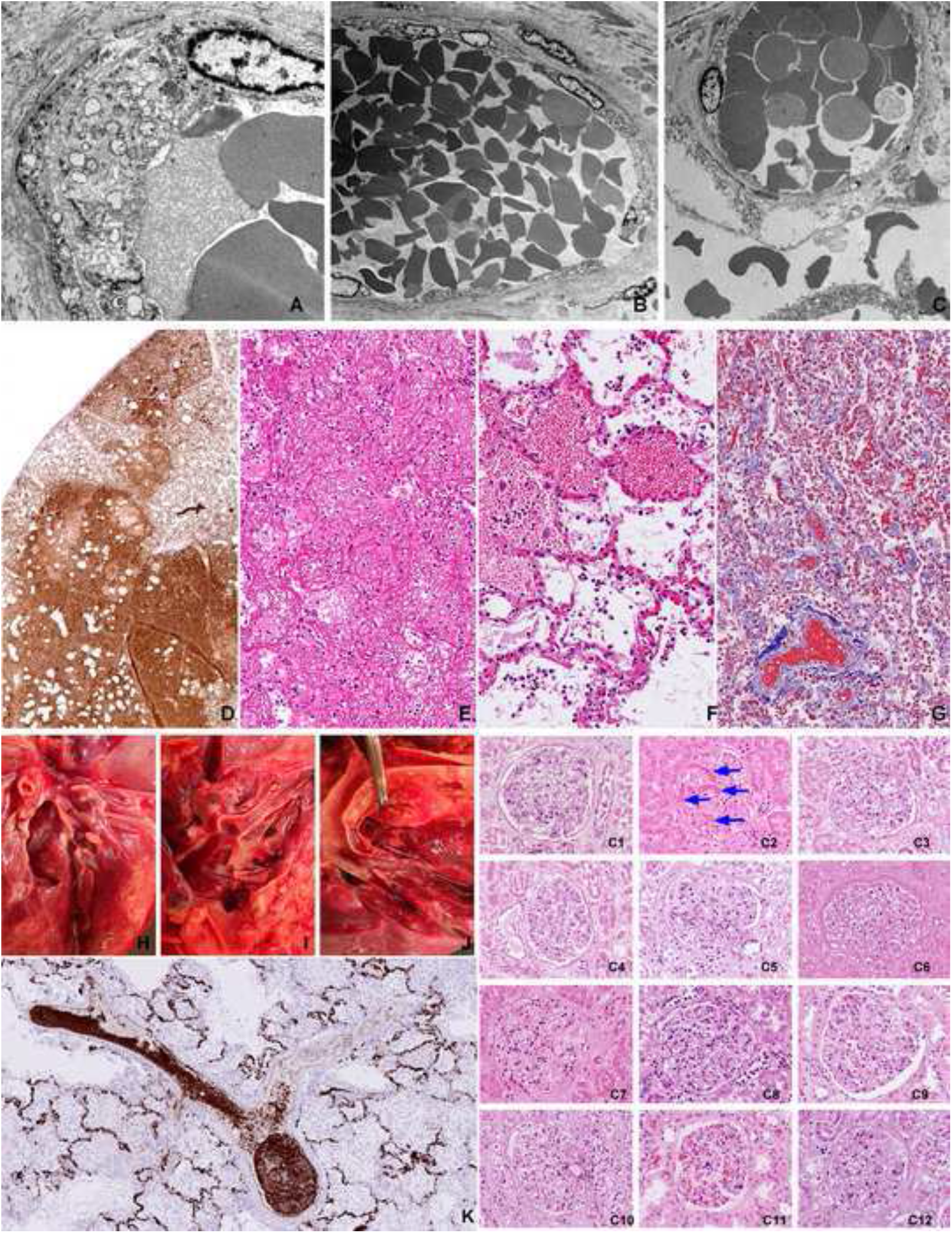
thrombo-embolic complications are rarely observed. Vacuolising cytopathic effect on the capillary endothelial cells (A) that leads to fenestration (B), plasma leakage and bleeding (C) into the alveolar space (top row TEM images). The extent of plasma leakage is conveniently demonstrated by staining for von Vilebrand factor in the affected lung (D). Coagulated plasma in the alveoli constitute the largest amount of filamentous fibrin in the body of the Covid-19 victims (E) HE staining. Intra-alveolar bleeding from alveolar wall capillaries (F) HE staining. Collapsed alveolar walls in progressing consolidation where the alveolar lumens are filled with coagulated blood (Masson trichrome). Thromboembolic complications were uncommon in our cohort. The multiple thrombemboli that was observed in C8 (left panel upper row) was mainly composed of aggregated CD61 positive thrombocytes (K) rather than fibrin coagulum (left panel bottom). Disseminatied intravascular coagulation was observed only in one case (C2) as shown in the glomerulus of the kidney (right panel).

The most frequently mentioned pathological change in relation to Covid-19 is the description of thromboembolic complications especially the detection of fibrin micro-thrombi. In our autopsy series these structures were rarely observed. One victim (C8) had extensive intra-vascular thrombus formations in the branches of pulmonary arteries. These thrombi however were mainly composed of aggregated thrombocytes as was demonstrated by staining with CD61 and CD31 (Figure 3 K). Despite of massive accumulation of thrombocytes in the pulmonary intravascular lumen, there were very few thrombocytes in the intra-alveolar bleedings or in fibrin coagulums. Megakaryocytes were frequently observed in the pulmonary vasculature in most cases.

Only one patient (C2) suffered from disseminated intravascular coagulation (DIC) as judged by analyzing kidney glomeruli for the presence of fibrin micro-thrombi (Figure 3 C1-C11). Most pulmonary vessels were free of thrombi even in this patient.

Recruitment of inflammatory cells is a salient feature of most infectious conditions. Virus induced pneumonias are often aggravated by bacterial superinfections. In the current cohort we observed only one case of additional purulent trachea-bronchitis/pneumonia. Neutrophil granulocytic infiltrations were regularly absent from the lungs of this cohort. The pulmonary inflammatory infiltrate contained slightly elevated levels of CD2, CD3, CD4 positive T cells with an almost complete absence of CD8 positive T killer cells, CD56 positive NK cells and CD20 positive B cells. In line with these data there were only a few perforin and granzyme B positive cytotoxic effector cells that only occasionally engaged with the virus infected cells. The mononuclear inflammatory infiltrate was regularly dominated by myeloid cells with great cell size variation ranging from small myeloblast-like cells with narrow cytoplasmic rim to multinucleated giant cells. These myeloid cells were regularly positive for CD14, CD16, CD31, CD33, CD43, CD45, CD68, CD117 and CD163. These cells also expressed tartrate resistant acid phosphatase (TRAP), alfa1-antitrypsin and high levels of heatshock protein Hsp70. The surface of these myeloid cells was coated with von Willebrand factor, IgM and C reactive protein. The myeloid cells constituted a large fraction of the nuclear cells of the organized consolidations reaching 50-80% of all nucleated cells in many areas. Many of the myeloid cells were actively proliferating as shown by positivity for Ki67 and phospho-Histone H3. The mediastinal and hilar lymph nodes were regularly enlarged but there was no generalized lymphadenopathy at other sites. The enlarged hilar nodes showed recognizable germinal center structure but with the absence of centroblasts. The GC cells had maintained expression of CD20, CD23 but greatly decreased expression of CD10. There was a marked reduction of Ki67 in the germinal centers. There was a great overall reduction of CD8 and CD57 positive T cells. There was a great increase of Ki67 positive proliferating myeloid cells in the medulla that were also CD163 and TRAP positives and were coated with von Willebrand factor and IgM.

We have observed diffuse alveolar damage in all Covid-19 victims with massive production of hyaline membrane structures. Immunohistochemical analysis of hyaline membranes revealed that they contained remnants of the debris of alveolar epithelial cells (CK18, EMA, Muc1, Muc5AC, surfactant), macrophages (CD68, CD14, CD163, CD117, CD31, vimentin) and serum proteins (IgM, CRP, von Willebrand factor) (Figure 4). We observed ample deposition of virus RNA positive cell debris from infected cells in the hyaline membrane fragments.

**Figure 4.**
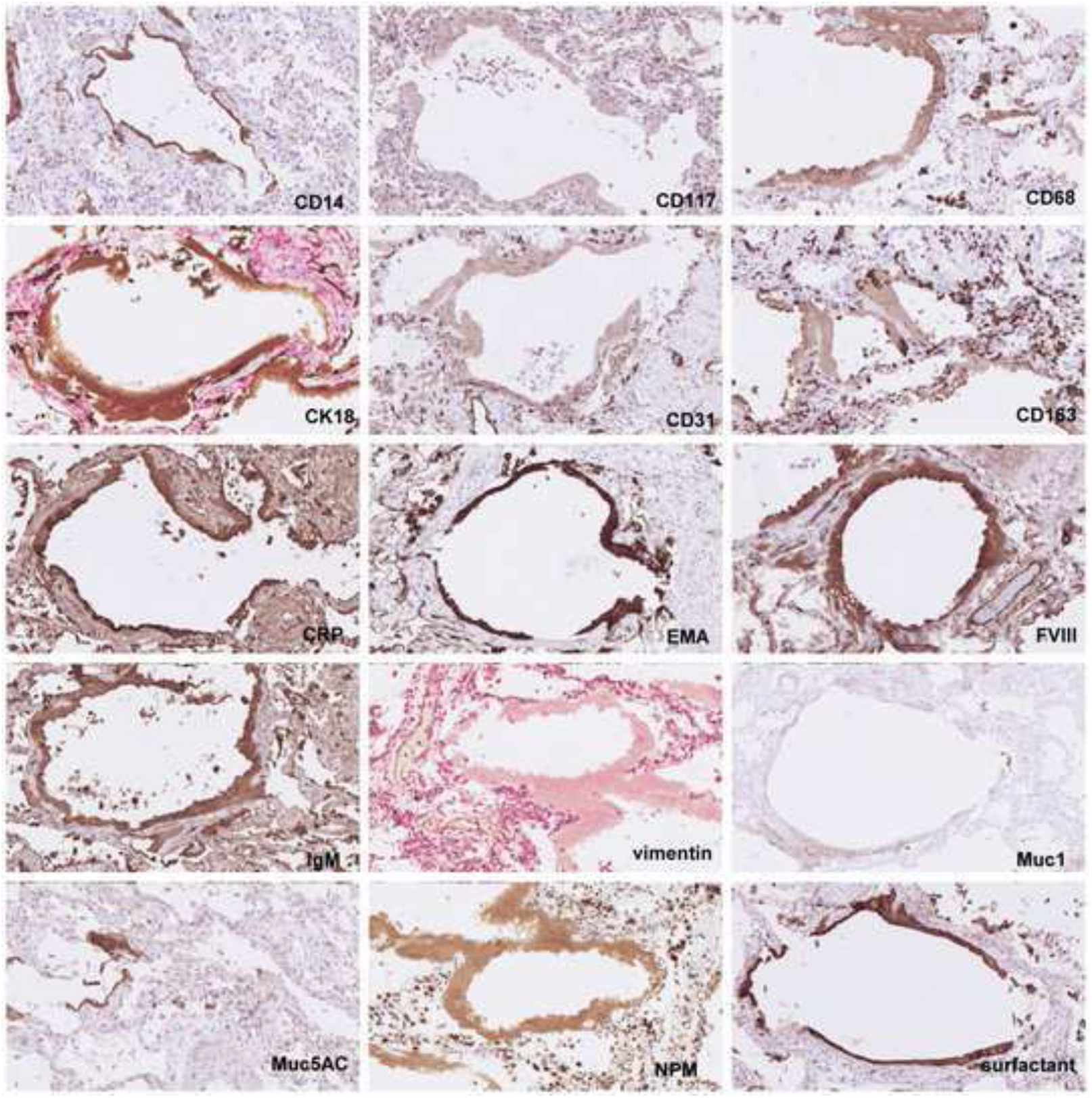
Identification of hyaline membranes. Molecular composition of the hyaline membranes reveal that they are composed of cell debris of epithelial cells, macrophages and serum components.

The process of lung consolidation was accompanied with massive accumulation of macrophages and immature myeloid elements, extensive proliferative response both in the epithelial and stromal components as well as stromal reaction with the emergence of podofillin and alfa-smooth muscle actin positive stromal cells and exuberant neo-angiogenesis (Figure 5).

**Figure 5.**
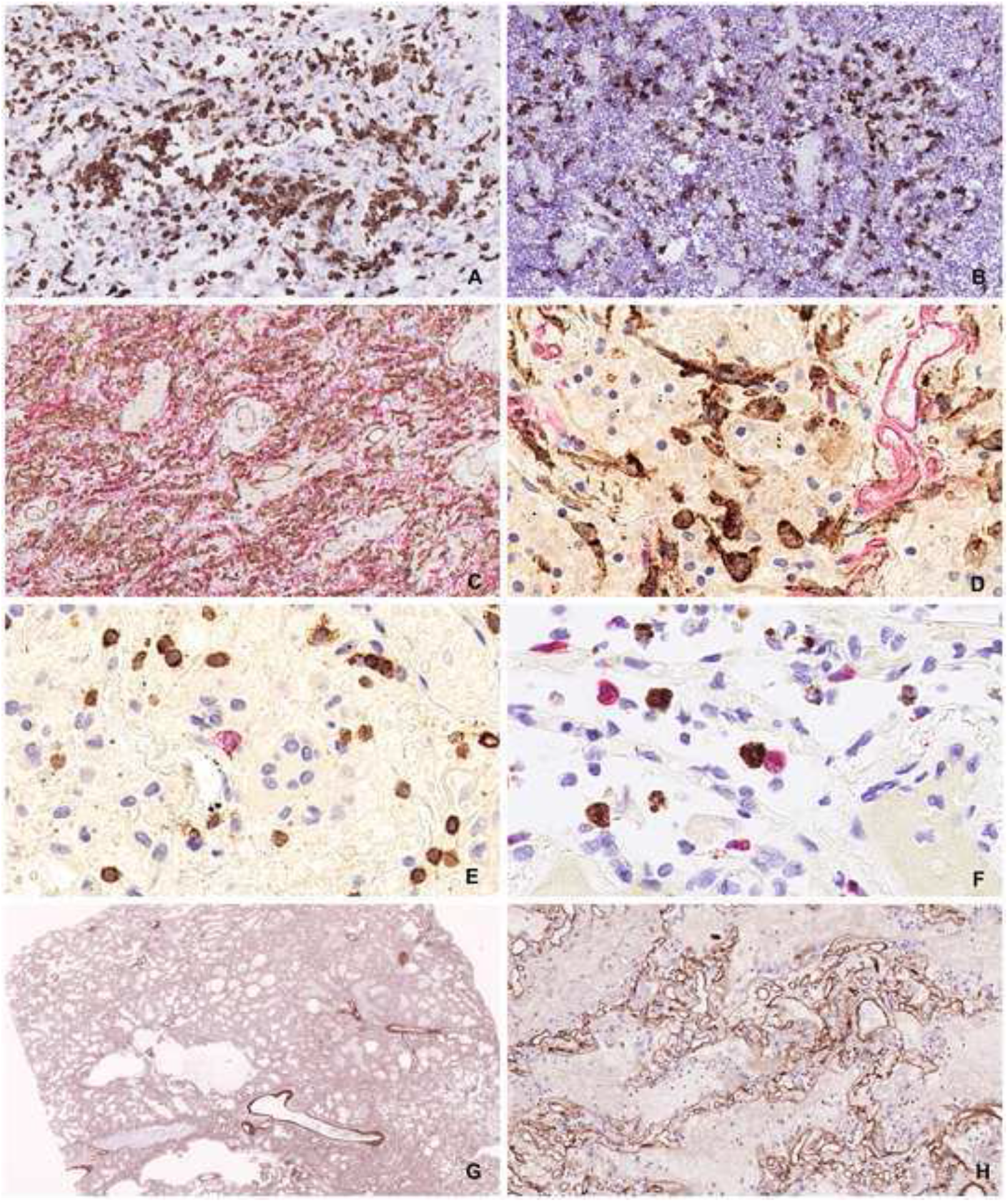
Cellular events of pulmonary consolidation reaction. Accumulation of large number of CD163 positive myeloid cells in the lung parenchyma (A) and in the hilar lymph nodes (B) as a sign of MAS (macrophage activation syndrome). Robust angiogenesis with the production of large number of new capillaries with patent lumen (C) Proliferation of podifyllin (D2-40 positive stromal cells (D). Subdued lymphocytic reaction with decreased amount of B cells (CD29 – red) and modest amount of T cells (CD3-brown) (E). Rare event of an immune engagement of a virus infected cell (RNAScope brown) with a Granzyme B positive effector cells (F). Preservation of the collagen IV scaffold inside the heavily consolidated area might provide mechanical framework for later tissue regeneration after clearing off the solid elements (G and H)

#### Circulatory system

The atherosclerotic involvement of the aorta, coronary arteries, the kidney and the basal artery is summarized in Table III. Only two patients (C7 and C10) showed advanced atherosclerotic lesions in all vessels examined. The majority of the patients (9/12) showed hypertrophy of the wall of the left cardiac chamber, four of them even had right chamber hypertrophy. Dilatation of the right chamber was observed in nine cases. One patient showed the signs of old healed myocardial infarction in the posterior wall of the left chamber (C7). Three patients showed very extensive hypoxic damage of the entire myocardium in the form of contract band necrosis at multiple sites (Supplemental Figure 3.)

### CNS

All brains were perfused and sliced in a series of coronal sections. Brain edema was the most prominent pathological change (6/12). No major bleeding, macroscopically notable inflammation or malacia was observable. Occasional dilatation of blood vessels was noted. Histologically perivascular edema and focal perivascular bleeding could be detected along with minimal cytopathic changes of neurons in the basal ganglia (Supplemental Figure 4).

### Gastro-intestinal system

No systemic macroscopic pathological change was observed in the GI tract. Microscopic examination of the mucous membranes revealed extensive denudation of intestinal villi even in the rapid autopsy cases, a phenomenon that was considered to be the result of the generalized pre-mortem hypoxemia.

### Detection of the virus in various organs

The reliability and sensitivity of different in situ virus detection methods were tested on infected lung tissues from rapid autopsies (C9 and c11). RNAScope and immunohistochemistry using antibodies against Sars-Cov2 nucleoprotein and Spike protein gave very similar distribution of the virus infected cells. RNAScope was far the most sensitive method. Systematic testing of all organs from a male and female donor from rapid autopsy (C9 and C11) revealed that the greatest amount of virus infected cells was detected in the lung alveoli with minimum change (edema, incipient bleeding, epithelium slough-off and proliferation) (Figure 6). Remnants of virus infected cells with spread out lytic cell debris were detectable in the areas of early consolidations (fibrin coagulum, atelectasia, macrophage accumulation). Virus was regularly absent from areas of late stage consolidation (stromal reaction, neoangiogenes). No viral replication was detected in the epithelium of the tongue root, pharynx, epiglottis, vocal cord, trachea, large and small bronchi although ample virus material was visible in the lumen or smeared over the surface of the epithelia. Systematic testing of all organs yielded rare, scattered, single cells in various organs such as brain, heart, intestines, kidney, rectum, thyroid, spleen, no evidence for foci of virus replication was found. We concluded that the main site of virus replication was the alveolar space of the lung parenchyma.

**Figure 6.**
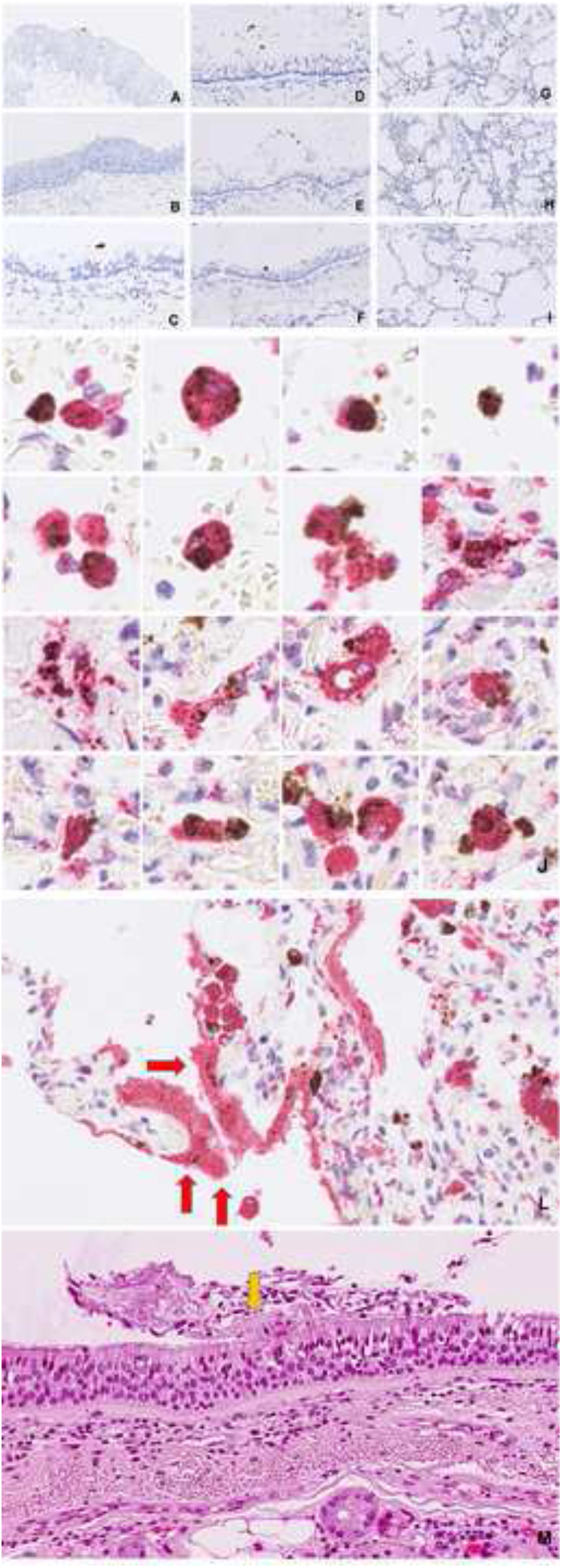
Distribution of virus infected cells in the airways. Virus RNA positive material is restricted to the lumens of pharynx (A), epiglottis (B), trachea (C), man bronchus (D) and minor bronchi (E,F) whereas there was ample virus signal carrying cells in the alveoli (G,H,I). Productive virus infection of the alveolar macrophages is demonstrated by combination of RNAScope detection of viral Spike RNA and staining of CD68 macrophage marker (J). Deposition of virus carrying debris in the hyaline membranes (L). Hyaline membrane fragment on the surface of the trachea (M)

### Isolation of the Sars-Cov2 from the lung tissues

Cell free, 0.22 um filtered fluid of homogenized lung tissue of case C9 was incubated with VeroE6 monolayers and observed for cytopathic effects. The presence of the virus in the infected cell cultures was demonstrated with RT-PCR. The viral isolate, Sars-Cov2-KH78 showed vigorous replication inducing mild cytopathic effect after 24 hours, severe cytopathic effect after 48 hours and the death and lysis of >95% of the target cells after 72 hours (Figure 7).

**Figure 7.**
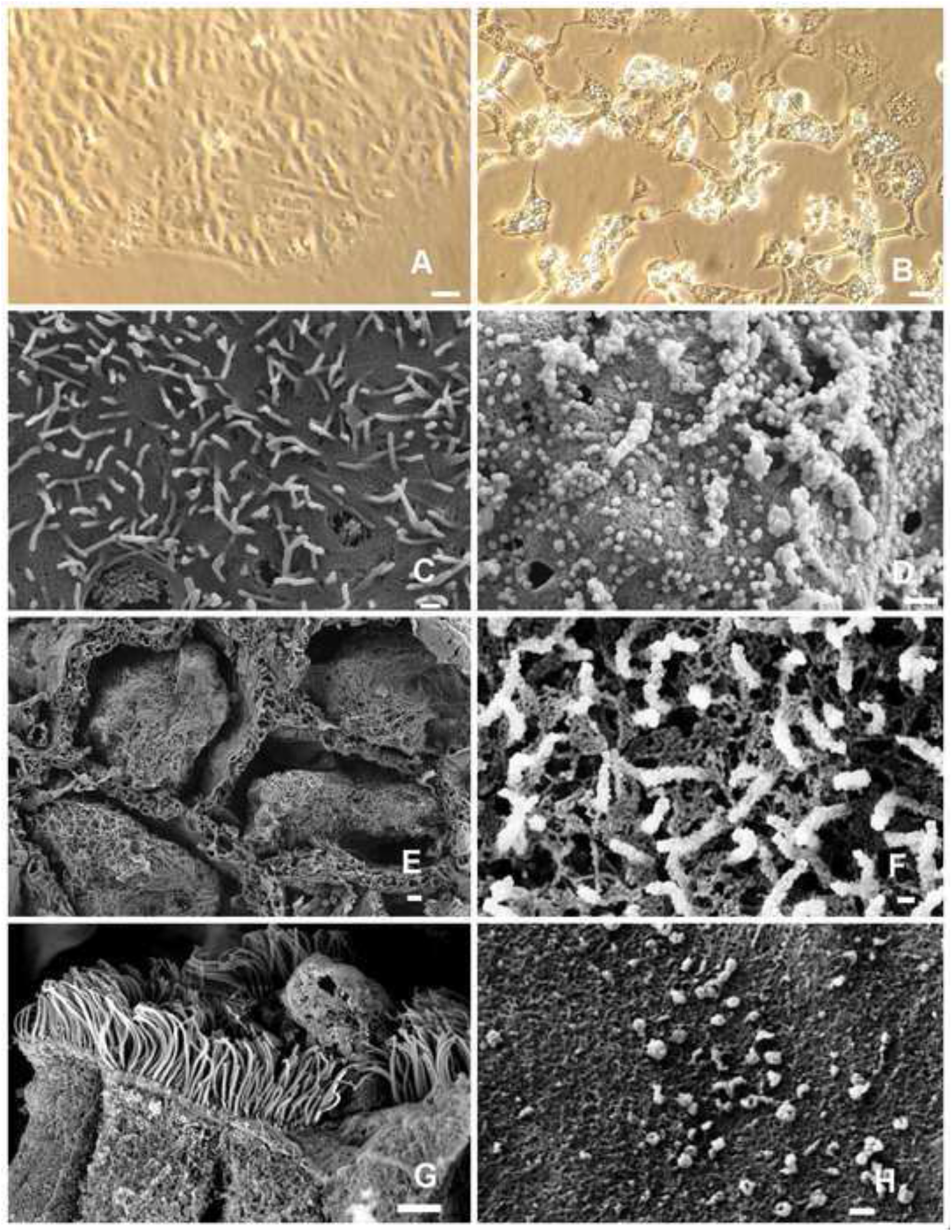
Vitro study of isolated virus. Isolation of the virus from the lung of C9 using Vero E6 cells (A - uninfected). Cypathic effect on the VeroE6 48 hours after infection (B). Phase contrast (A and B) Scanning electron microscopy (SEM) images of the surface of uninfected VeroE6 cells (C) and infected Vero E6 cells 48 hours post infection (D). Intra-alveolar coagulum (E). Budding virus particles on the surface of a type II pneumocyte. Virus infected dead cell in the bronchiolar lumen, on the top of ciliated epithelium (G) Virus particles on the alveolus wall (G). E,F,G and H SEM images from the lung of patient C9. Size markers A and B 10 micrometer, C and D 300 nanometer, E 10 micrometer, F 100 nanometer, G 2 micrometer, H 100 nanometer.

## Discussion

Adequate therapy of a virus infection initiated sickness require the proper understanding of not only the virus infection induced cell damage but also the effect of different host responses. As most victims of Covid-19 are in advanced age with fragile health and numerous co-morbidities, the delineation of the virus induced pathologies require patient cohorts where the major tissue damage and eventually the direct cause of death can be directly linked to the presence of the replicating virus. Sars-Cov2 is considered a respiratory virus with lung damage as leading pathology. Our cohort comprises individuals who succumbed to the devastating effects of virus initiated lung damage, the Covid-19 associated acute respiratory distress syndrome.

The gross changes of the lung parenchyma was similar to cases reported by others (Suess & Hausmann, 2020)(Bradley et al., 2020)(Zhou et al., 2020)(Barton et al., 2020)(Martines et al., 2020)(Adachi et al., 2020). In this cohort however we found that most macroscopic and microscopic pathologies were observed in the lung parenchyma as opposed to the upper or lower airways.

Our findings suggest that in these patients the main site of the virus replication is in the pulmonary alveoli and not the laryngeal, pharyngeal, tracheal or bronchus epithelia. The leading pathology that eventually destroys the lung respiratory functions appears to be the massive consolidation of the lung parenchyma. The process is initiated by virus replication in the pneumocytes leading to the desquamation of the alveolar epithelia and various levels of breach in the barriers between alveolar capillaries and the intra-alveolar space. With the increasing severity of this breach edema fluid, plasma and eventually whole blood is leaking out into the alveolar space. Initiation of the coagulation cascade leads to the accumulation of fibrin filaments and/or intra-alveolar coagulated blood. Importantly only a minority of peumocytes showed lytic virus replication but most pneumocytes in the affected area showed pronounced cytopathic effects in form of cytoplasmic swelling, vesicular degeneration and nuclear atypia. At the present it is unclear how much of the epithelial damage can be interpreted as a by-stander effect or as a sign of abortive infection with arrested virus replication. Systematic mapping of early viral proteins such as the intact or cleaved products of the ORF1a and 1b would be required to answer this question.

It has been suggested that various cellular elements of the vasculature such as endothelia (Varga et al., 2020) or pericytes (He et al., 2020) could be direct targets of Sars-Cov2 infection or would be damaged by subsequent inflammatory cell infiltrates (Becker, 2020). We have never seen any sign of viral presence in any of these structures nor inflammatory reaction against vascular wall components in our cohort. On the other hand we clearly demonstrate various levels of cytopathic effects of endothelial cells in the vicinity of virus infected epithelial cells by immunohistochemistry and electron microscopy. We also show that local release of red blood cells into the alveolar space is linked to the presence of virus infected epithelial cells. We interpret the endothelial damage as a by-stander effect possibly caused by soluble factors released from the infected pneumocytes. One candidate for such an effect might be the product of ORF3a, a viropore protein that has been shown to induce apoptosis in non-infected cells in SARS (C. M. Chan et al., 2009) and Covid-19(Ren et al., 2020).

Extensive neutrophil infiltration and NET formation was hypothesized to significantly contribute to the development of ARDS (Barnes et al., 2020). In our series neutrophils were rarely present in the lethal ARDS lesions.

All victims had high pre-mortem D-dimer levels in the serum similarly as reported in other studies (Mucha et al., 2020). High D-dimer together with well circumscribed radiological lesions regularly raises the clinical diagnosis of embolization of lung arteries. This diagnostic hypothesis was seemingly supported by pathology reports describing widespread intravascular fibrin micro thrombi implying generalized coagulopathy in Covid-19 patients (Calabrese et al., 2020)(Bradley et al., 2020)(Ackermann et al., 2020)(Vasquez-Bonilla et al., 2020)(Wichmann et al., 2020). In our series we had only one case with disseminated intravascular coagulation. One other case with numerous pulmonary arterial thromb-embolization was shown to be composed mostly of platelet thrombi. Our data is congruent with the notion of the presence of extensive coagulation events. We suggest however that in many severe cases the bulk of the coagulum may be situated in the intra-alveolar rather than in the intravascular space. The increased frequency of CD61 positive intrapulmonary megakaryocytes along with the numerous thrombocyte micro-trombi raises the question if thrombocyte aggregation inhibitor therapy might be more beneficial for advanced Covid-19 patients than the routinely employed anticoagulation therapy. The reason of intra-alveolar bleeding appears to be linked to ill-defined cytopathic effects on vascular endothel in the vicinity of virus infected cells.

It has been suggested that many of the pathological events of severe Covid-19 disease could be explained by virus induced cytokine storm characterized by increased levels of TNFalfa, IL-6, IL-8, IL-1beta and subsequent damage to the vascular endothelium (Huang et al., 2020). Recent data on severe Covid-19 ARDS cases showed much less pronounced cytokine elevations as compared to septic shock associated ARDS (Kox et al., 2020). Our data showing a great accumulation of CD163 positive myeloid cells in the lung consolidations and hilar lymph nodes in parallel with CD8 T, NK and B cell depletions in these tissues are in line with previous suggestions of M2 polarized macrophage induced effector T cell depletion (Liu et al., 2020)(Schulte-Schrepping et al., 2020). The elevated level of CD163 with the subsequent shedding is an emerging marker of virus infection associated macrophage activation syndrome with dismal outcome (McElroy et al., 2019) (Loomba et al., 2020).

The lung parenchymal consolidation was marked by a vigorous proliferation activity in the lung epitelium, endothelium, stroma and myelo-monocytic compartment. All these activities appear to compromise normal respiratory functions thus general proliferation inhibition with cytostatic chemicals or radiation might have a therapeutic value. Similarly, inhibition of the vigorous neoangiogenesis with anti-VEGF antibodies might be justified.

The presence of the virus was reported in several tissues in various studies, mainly based on RT-PCR based detection (Hanley et al., 2020) or replication competence in ex vivo cultures (Hui et al., 2020). Already the earliest studies described of “viral RNAmy” in the blood of some of the virus carriers (Huang et al., 2020). Our systematic mapping revealed a number of scattered cells in most organs without direct evidence of local virus replication outside of the alveolar space.

Most cellular RNA degrades after 48 hours postmortem thus making virus detection in infected cells by RNA in situ hybridization very difficult. The virus however can be detected from infected lungs up to twelve days by RT-PCR (Edler et al., 2020) suggesting that the viral RNA persists well protected in the viral capsids. Our data show that extensive deposition of virus RNA in the hyaline membranes raises the possibility that released viral particles may be embedded in large quantities in the protective matrix of the hyaline membrane. Detection of hyaline membrane fragments in the upper airways may provide an alternative route for viral spreading. We suggest that the intra-alveolarly produced virus might leave the body with the exhaled air deposited in the proteinaceous matrix of the hyaline membrane. This matrix may provide extra environmental protection and possibly prolong the survival of the virus ex vivo. Both Sars-Cov (K. H. Chan et al., 2011) and Sars-Cov2 (Doremalen, Bushmaker, 2020) may remain viable on various surfaces for days when sprayed out as cell culture supernatant. Virus entrapped in dried out protein matrix may be prevented with receptor expressing target cells unless it is actively released. Inhaled virus bearing hyaline membrane particles are likely ingested by alveolar macrophages where the acidic environment and protease activity in the endosomes might initiate the generation of Spike fusion complex. Our data show that the frequent presence of viral RNA in CD68 positive macrophages is congruent with the hypothesis that phagocytic cells consuming virus carrying cellular debris might themselves become infected. This would provide an alveolus to alveolus transmission route through aerosol spreading that does not require replication events in ACE2 expressing cells. The absence of replicating virus in the upper airways and the focal, patchy, mosaic like involvement of the terminal lobules is also consistent with this type of viral spread.

Alternative way for Sars-Cov2 entry into macrophages is through antiviral antibodies that may function as adapters through Fc receptor mediated viral binding and receptor mediated endocytosis. Macrophage infection through Fc receptors can lead to antibody dependent enhancement of the disease (ADE) that is well documented in vaccination models of dengue (Srikiatkhachorn & Yoon, 2016), feline infectious peritonitis (Takano et al., 2008)(Kipar & Meli, 2014) and experimental Sars vaccine models (Luo et al., 2018). The argument that Covid-19 vaccine would be free of ADE is largely based on the data that macrophages are refractory to Sars-Cov2 infection in vitro when cultured virus is used as infective agent. The demonstration of Spike RNA in CD68 positive cells in the infected lungs however raises the concern that certain vaccinated individuals who produce neutralizing antibodies against the Spike protein could succumb to ADE upon exposition to the circulating virus if the antibody would function as an adapter to bring the virus into the macrophages (Arvin et al., 2020)(Peeples, 2020)(Eroshenko et al., 2020). In vitro macrophage infection experiments using antibodies from victims who succumbed to Covid-19 will be required to test the validity of this scenario.

The added value of this study, beside detailed characterization of the lethal pulmonary injury caused by the virus, is the precise localization of the virus infected cells using RNA in situ hybridization in relation specific cell types identified by to immunohistochemical markers. Our data indicate massive bystander effect beyond the direct virus induced cytopathic damage. We show endothelium damage in the vicinity of virus replication without the presence of the virus in the endothelium. We present evidence of macrophage infection in the lungs. This finding can have major effect on the evaluation of the risk of ADE. We localize most of the virus production in the alveolar space and not in the upper airways, raising the possibility for alveolus to alveolus spread of the virus possibly embedded in hyaline membrane material. We show that in most fatal cases the fibrin coagulation happens in the intra-alveolar space and not in the vessels. We also show that thromboembolic events might be caused by increased thrombocyte aggregation rather than by increased intravascular coagulation. We show that massive proliferation activity in the epithelial, stromal, vascular and myelo-monocytic compartment contributes to the development of lung consolidations. Our data suggest that thrombocyte aggregation inhibition, angiogenesis inhibition and general proliferation inhibition may have a roll in the treatment of advanced Covid-19 ARDS.

## Acknowledgements

This study was based on clinical autopsies financed by region Stockholm, Karolinska University Hospital.

## Authors contributions

BSL3 autopsies were carried out by LS and ASz. Grossing was done by LS, YW and XY. Histological and immunohistochemical analysis and interpretation was done by LS, BB, MB, YC, MBj, ASz, immunohistochemistry and RNAScope stainings were done by MO and PL, virus isolation and culturing was done by LS, SG, ES and AS, electronmicroscopy was done by LH and LS. Manuscript was written by LS, BB, MB, YC, MBj, AS and ASz. Figures were assembled by LS, BB, YW, XY and ASz. All authors red and approved the manuscript.

## STAR Methods

### BSL-3 level autopsy

All autopsies were conducted at the risk-autopsy facility of the Department of Clinical Pathology/Cytology, Karolinska University Hospital, Huddinge, Stockholm, Sweden. All subjects were referred to the pathology department for clinical autopsy to establish the precise cause of death. The study was approved by the Swedish Ethical Review Authority under the registration number DNR 2020-02446 and 2020-04339.

All postmortems were carried out by a two man team of clinical pathology specialists (L.S and A.S) using a COVID-19 adapted “autopsy in-the-bag” procedure that allows the detailed examination, photo documentation, histological, virological and electron microscopic sampling of all organs without the use of running water and without the production of infectious aerosols. Briefly the bodies were opened through a Y shaped skin incision from the shoulders to the symphysis. The anterior wall of thorax was removed and the abdominal cavity was opened. Live samples from various organs were taken under aseptic conditions for virological analysis. Upon removal of the heart the two carotid arteries and subclavian arteries were cannulated and the head and neck area was embalmed to prevent the generation of infectious aerosols when using the oscillating autopsy saw during the brain removal. The organs were removed in a single block starting from the tongue and soft palate all the way down to the rectum. Subsequently the thorax organs were separated from the abdominal organs. All regions were inspected and photographed and ample amount of histological samples were collected for further analysis. The following organs were routinely sampled: soft palate, tongue root, pharynx, larynx, epiglottis, vocal cord, trachea, main bronchus, small bronchi, mediastinal lymph nodes, multiple sites of lung parenchyma, heart, esophagus, jejunum, stomach, sigmoid, rectum, liver, pancreas, kidney, bladder, spleen, adrenals, thyroid, testis, prostate, brain. All subsequent procedures were carried out in at our clinical laboratory accredited to issue medical diagnosis.

### Processing of histology samples

Samples of all relevant organs were collected in 4% buffered formaldehyde were subsequently excised for paraffin embedding and sectioning after minimum three days of fixation. Forty to sixty tissue blocks were generated from every patient for routine histology. The organ fragments were repeatedly photographed before and after grossing to ascertain the exact position of the tissue blocks in relation to relevant anatomical structures. The blocks were de-hydrated and paraffin embedded using automated tissue processors. Five micrometer sections were cut from the paraffin blocks and routinely stained hematoxylin-eosin in staining automated devices.

### Immunohistochemistry

Immunostaining was performed in Ventana Ultra Benchmark (Ventana Medical Systems, Inc., 1910 Innovation Park Drive Tucson, Arizona USA) according the manufacturer’s instructions.

For the complete list of antibodies see supplementary table 1.

RNA in situ hybridization

### RNAscope^®^ ISH Procedure

Human standard probes for use on automated Bond III Leica Systems were used from Advanced Cell Diagnostics (ACD). SARS-CoV-2 (bp 21631 – 23303, ACD # 848568), 4-hydroxy-tetrahydrodipicolinate reductase (dapB) (bp 414 - 862, ACD # 312038), was used as negative probe control. Ubiquitin C (UBC) (bp 342 – 1503, ACD # 312028) was used as positive probe control. The RNAscope target retrieval procedure “95” was chosen for the following tissues: heart, tongue, salivary gland, epiglottis, vocal cord, pharynx, trachea, lung, esophagus, bone marrow, liver, kidney, intestines, skeletal muscle, testis, urinary bladder, brain, adrenal gland, pancreas, stomach, uterus and cerebellum, according to the manufacturer’s instructions. The RNAscope target retrieval procedure “88” was chosen for lymph nodes, tonsil, thyroid and spleen.

### RNAscope^®^ ISH combined with IHC

In order to study different cells infected with SARS-CoV-2 virus RNAscope ISH was combined with conventional IHC staining with Alkaline Phosphatase Red using Bond Polymer Refine Red Detection Kit (DS9390). Directly after the ISH procedure the staining continued with IHC without further pretreatments. The antibody incubation time was doubled to 30 min from standard 15 minutes. Following antibodies were used: CD34 (clone QBEnd/10 form Dako # M7165, 1:25), CD68 (clone PG-M1 from Dako #M0876, 1:50), CK18 (clone DC-10 from Dako #M7010, 1:25), PDL1 (clone BSR90 from NordicBiosite # BSH-4003, 1:200), Granzyme B (clone 11F1 from Novocastra #NCL-L-GRAN-B, 1:25).

### Digital image processing

All slides were scanned at maximum (40x) resolution on Hamamatsu 360 Digital slide scanner and visualized by NDP Viewer. Over four thousand images in excess of 4.6 Terabyte were generated during this study.

### Virus isolation

One-gram unfixed piece of lung was removed from the right middle lobe through a small incision between the ribs and kept frozen at −20 °C in a sterile test tube until further processing at the BSL3 virus laboratory. Upon thawing, the tissue was minced in millimeter size fragments using a sterile scissor, mixed with 1 ml serum free IMDM medium and was homogenized using disposable Teflon homogenizing piston in an Eppendorf tube. The homogenate was centrifuged at 14K rpm in an Eppendorf centrifuge for 5 minutes and the cell free supernatant was placed on a VeroE6 monolayer for 60 minutes. Additional 10 ml IMDM supplemented with 10% fetal bovine serum was added and the infected culture was incubated for 48 hours. Supernatant of the infected culture was filtrated on a 0.45 um filter and was analyzed by RT-PCR to confirm the presence of the virus. The virus containing supernatant was used propagate the virus on VeroE6 monolayers.

### Electron microscopy

#### TEM

The cells were fixed in 2.5% glutaraldehyde in 0.1M phosphate buffer pH 7.4 at room temperature for 1h followed by storage at +4°C. Following the primary fixation the cells were rinsed with 0.1M phosphate buffer and Milli-Q^®^ water before postfixed in 2% osmium tetroxide in 0.1M phosphate buffer, pH 7.4 at 4°C for 2 hours. The cells were then stepwise ethanol dehydrated followed by stepwise acetone/LX-112 infiltration and finally embedded in LX-112 (Ladd Research). Ultrathin sections (approximately 60-80nm) were prepared using an EM UC 7 ultramicrotome (Leica Microsystems) and contrasted with uranyl acetate followed by Reynolds lead citrate and examined in a HT7700 transmission electron microscope (Hitachi High-Tech) operated at 100 kV. Digital images were acquired using a 2kx2k Veleta CCD camera (Olympus Soft Imaging Solutions GmbH).

#### SEM

Cells grown on Thermanox™ coverslips (Thermo Fischer Scientific) were fixed by immersion in 2.5% glutaraldehyde in 0.1M phosphate buffer, pH 7.4. The coverslips were rinsed with 0.1M phosphate buffer and Milli-Q^®^ water prior to stepwise ethanol dehydration and critical-point-drying using carbon dioxide in an EM CPD 030 (Leica Microsystems). The coverslips were mounted on alumina specimen stubs using carbon adhesive tabs and sputter coated with a thin layer of platinum using a Q150T ES (Quorum Technologies). SEM images were acquired using an Ultra 55 field emission scanning electron microscope (Carl Zeiss Microscopy GmbH) at 3kV and the SE2 detector.

## Supplemental figures

**Supplemental Figure 1.**
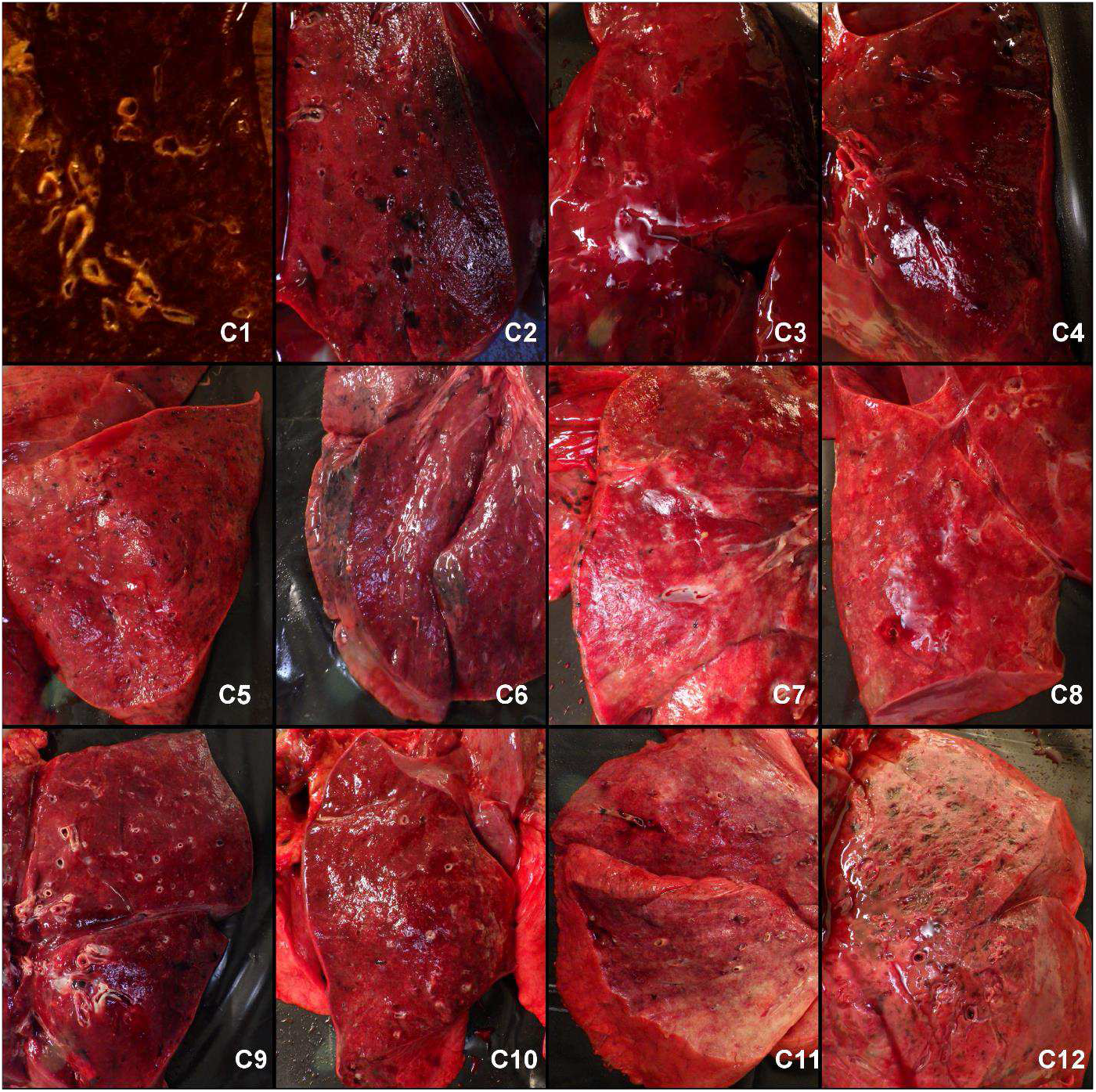
Cut surfaces of the lower lung lobes showed complete consolidation in all cases with total lack of air in the alveolar structures.

**Supplemental Figure 2.**
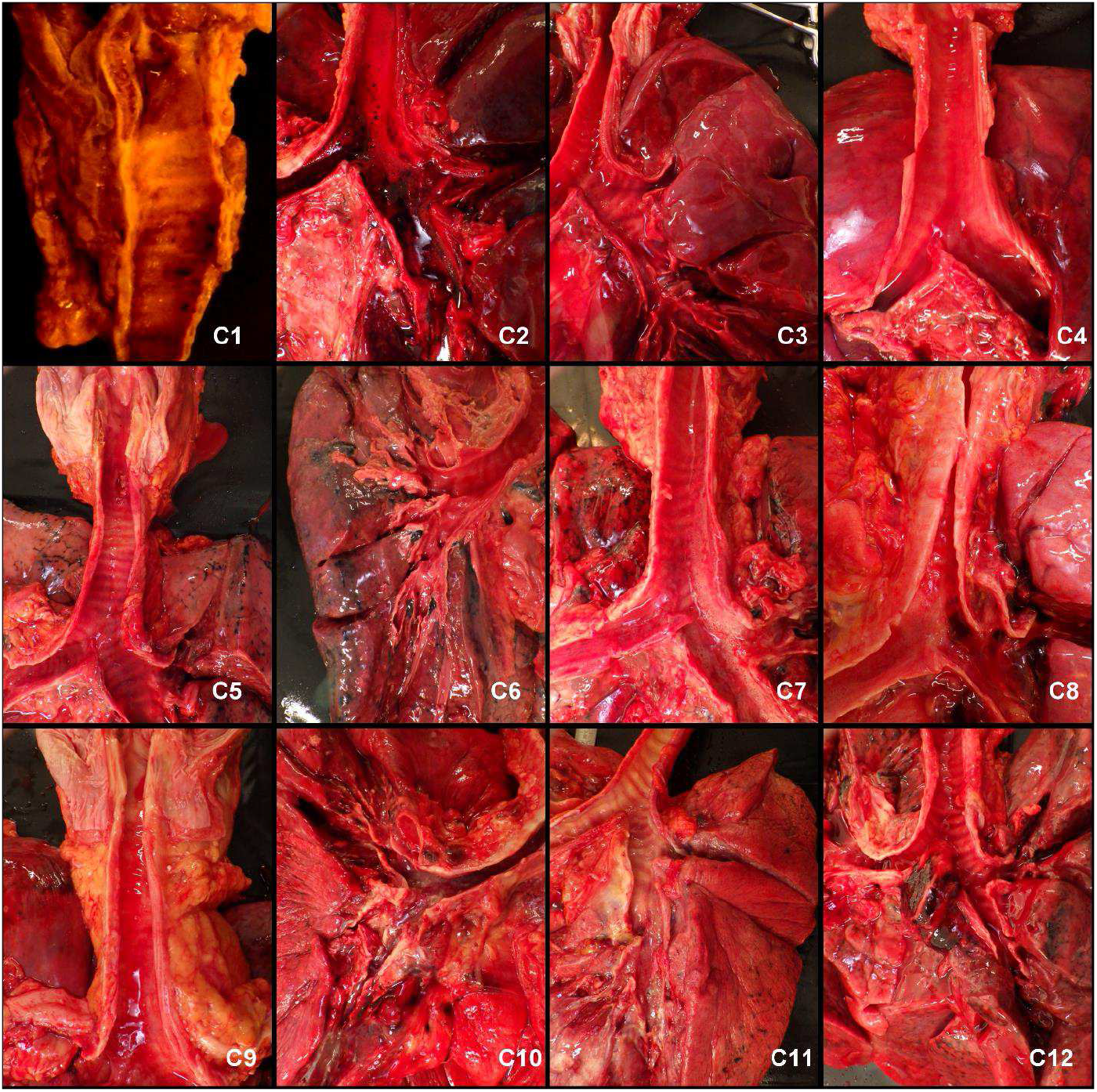
The upper airways were mostly unaffected.

**Supplemental Figure 3.**
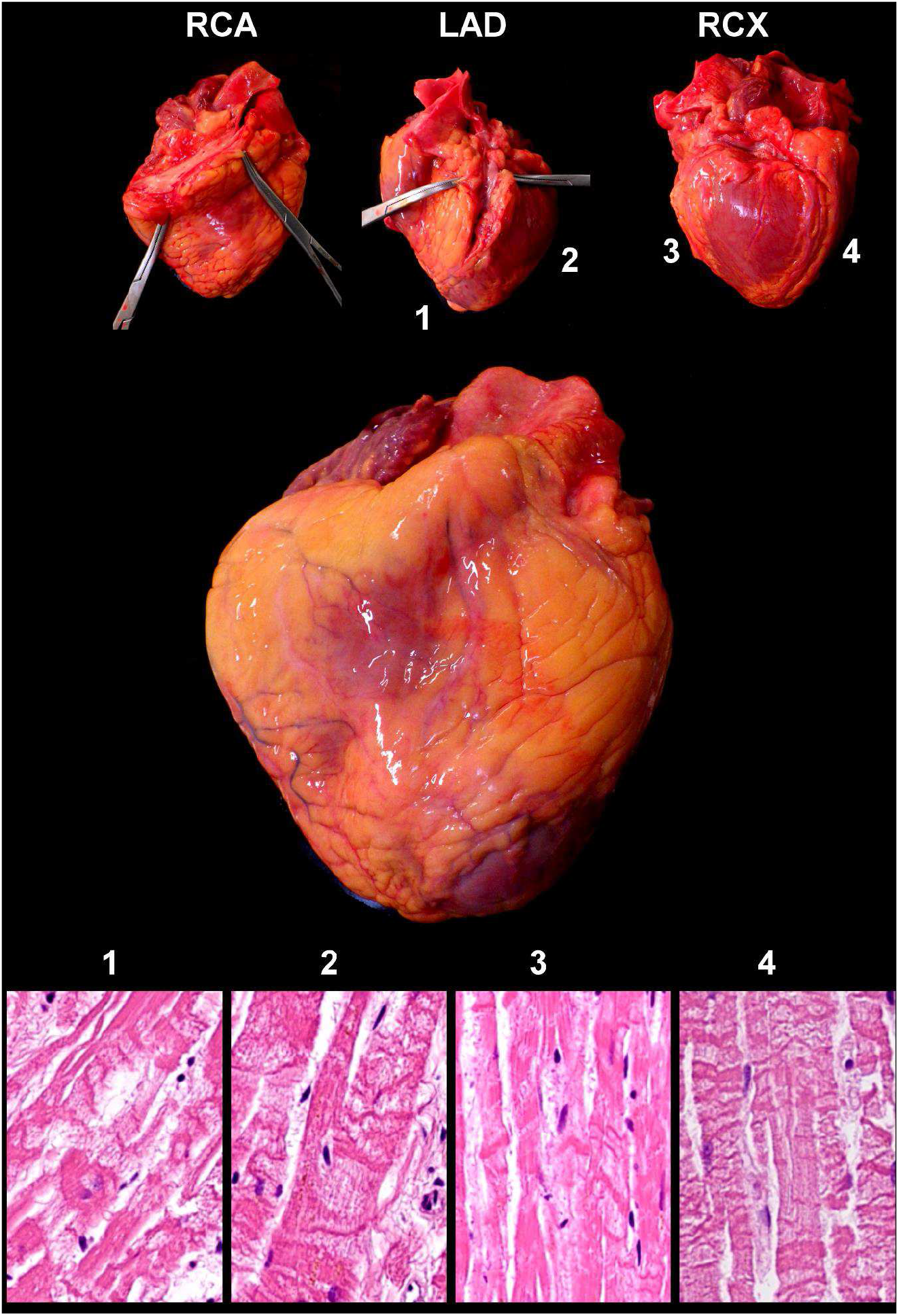
Generalized effect of tissue hypoxia on the heart of a Covid-19 victim. Massive contract band necrosis in all tested myocardium samples despite that all coronaries are fully open.

**Supplemental Figure 4.**
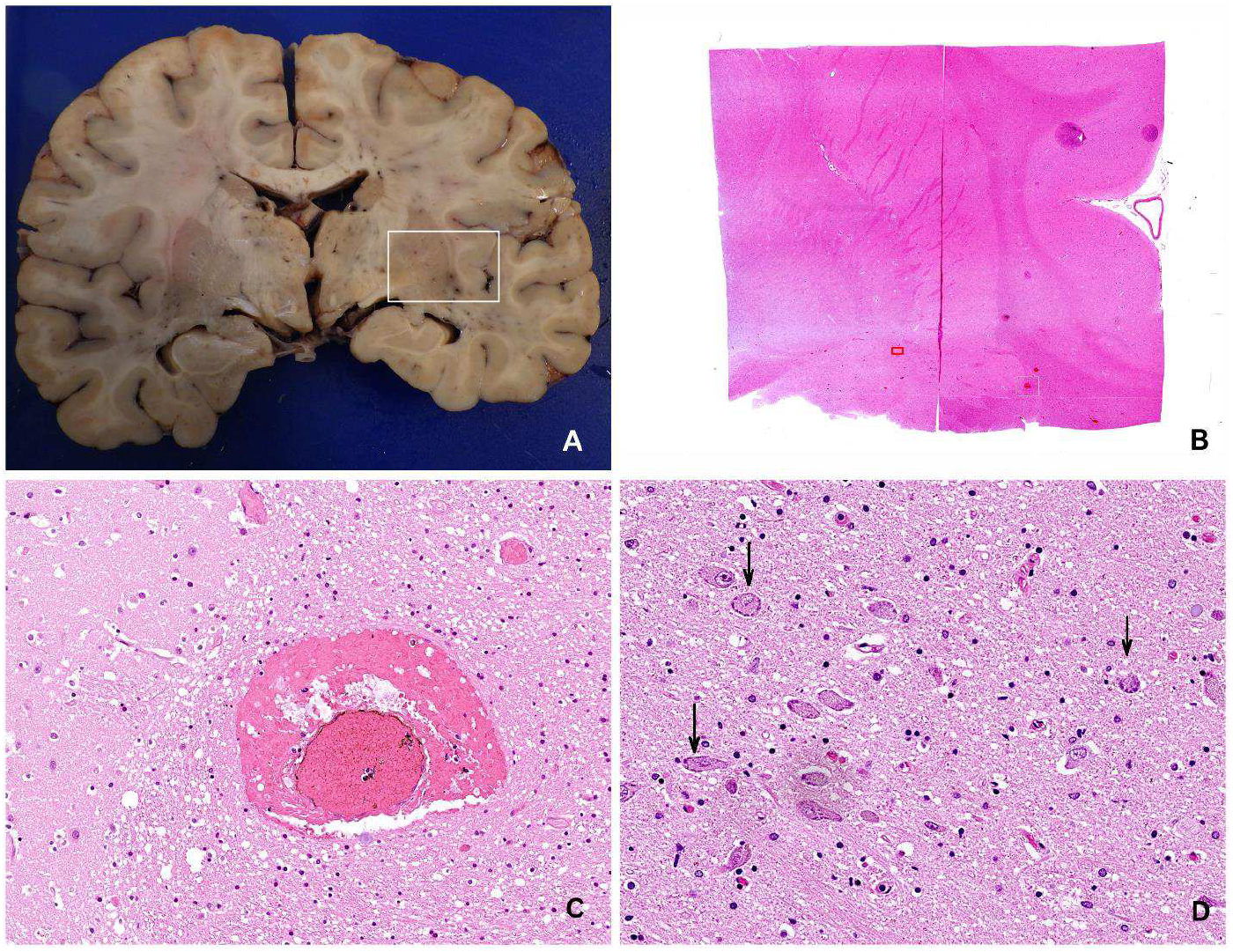
Minimal histological change in the most affected brain restricted to focal perivascular bleedings and subtle cytopathic effect in a few cortical neurons.

**Supplemental Table 1.** Patient characteristics

| Case (N) | Postmortem days | Days since the first symptom | Risk factors | BMI | D-dimer (<0.64) | CRP (<3) | Troponin(<15) | HLR | Ventillation | Died at |
| --- | --- | --- | --- | --- | --- | --- | --- | --- | --- | --- |
| 1 | 2 | 11 | DMII, liver transplatation immunosuppression | 31.9 | 29 | 153 | 606 | 10 min | 4days | ICU |
| 2 | 13 | 10 | no | 24.7 | ND | 199 | 428 | 45 min - LUCAS | no | home |
| 3 | 12 | 22 | no | 24.3 | ND | 327 | 22 | 55 min | 5 days | ICU |
| 4 | 11 | 24 | no | 33.1 | 3.6 | 152 | 21 | no | 7 days | ICU |
| 5 | 8 | 29 | DMII, Hypertonia | 33.7 | 27.3 | 247 | 276 | no | 21 days | ICU |
| 6 | 14 | 7 | DMII, Hypertonia | 24.2 | ND | 213 | ND | 1 hr- LUCAS | no | home |
| 7 | 5 | 11 | DMII, Hypertonia | 26.7 | 0.35 | 113 | 17 | 20 min | no | hospital ward |
| 8 | 3 | 14 | no | 18.7 | 35 | 147 | 92 | 90 min LUCAS | no | home |
| 9 | <1 (15 hours) | 7 | DMII, Hypertonia | 50.7 | >35 | 118 | 135 | 110 min | no | home |
| 10 | 5 | 24 | Hypertonia, stroke | 26.1 | ND | 256 | ND | no | no | hospital ward |
| 11 | <1 (9 hours) | 12 | chronic resp. insuff. | 26.9 | 2.8 | 250 | 80 | no | no | hospital ward |
| 12 | 7 | 30 | Hypertonia | 28.4 | 27.1 | 337 | 52 | 20 min | 7 days | ICU |
| **median** | 6 | 13 |  | 26.8 |  | 206 | 86 |  |  |  |

**Supplemental Table 2.** Organ weights

| Case | Height | Body Weight (kg) | Brain(g) | Heart(g) | Lungs(g) | Liver(g) | Kidney(g) | Spleen(g) |
| --- | --- | --- | --- | --- | --- | --- | --- | --- |
| 3 | 157 | 60 | 1310 | 290 | 1980 | 1680 | 310 | 120 |
| 4 | 179 | 106 | 1250 | 515 | 2570 | 2570 | 440 | 410 |
| 5 | 167 | 94 | 1259 | 395 | 2370 | 1570 | 410 | 410 |
| 6 | 170 | 70 | 1179 | 410 | 1250 | 1020 | 295 | 144 |
| 7 | 172 | 79 | 1369 | 390 | 1595 | 1440 | 295 | 310 |
| 8 | 176 | 58 | 1390 | 390 | 1630 | 1930 | 395 | 275 |
| 9 | 172 | 150 | 1540 | 790 | 2260 | 3160 | 460 | 655 |
| 10 | 175 | 80 | 1360 | 507 | 2370 | 1854 | 319 | 358 |
| 11 | 153 | 63 | 1160 | 554 | 1260 | 1997 | 277 | 128 |
| 12 | 179 | 91 | 1310 | 563 | 3654 | 2441 | 446 | 386 |

**Supplemental Table 3.**
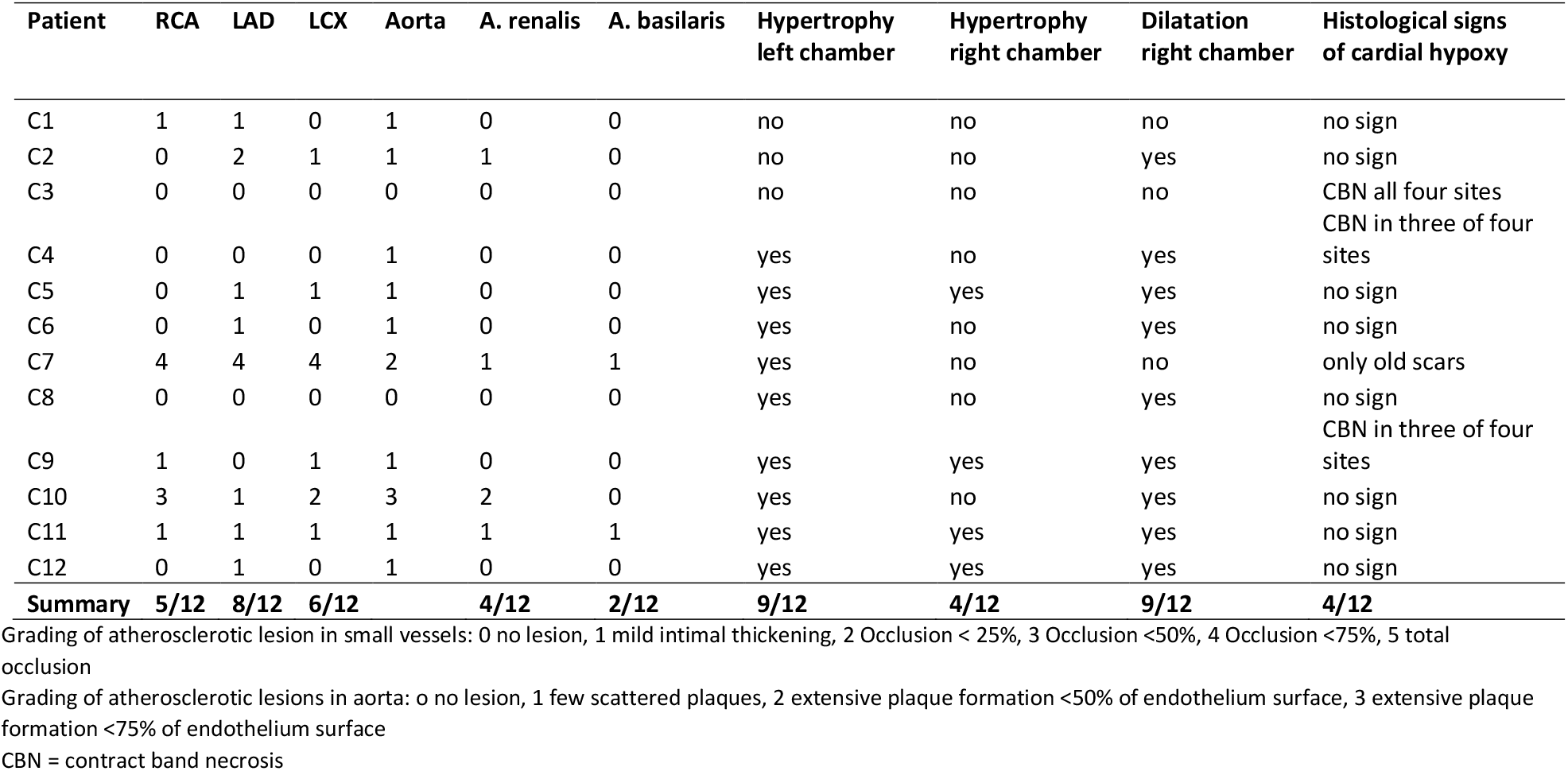
Cardiovascular pathology in Covid-19 victims

**Supplemental Table 4.**

| Primary antibody | Clone and catalog number | Company |
| --- | --- | --- |
| 5HMC | polyclonal # 39791 | Active Motif. |
| AAT | polyclonal # NCL-A1Ap | Novocastra. |
| ACE2 | clone 171606 # MAB933 | R & D Systems. |
| B-CAT | clone 14 # 05269016001 | Ventana. |
| CASP-3 | clone Asp175 # 9661 | Cell Signaling. |
| CD117 | clone 9.7 # 05278317001 | Ventana. |
| CD138 | clone B-A38 # 05269083001 | Ventana. |
| CD14 | Clone EPR3653 # 06419151001 | Ventana. |
| CD16 | clone 2H7 # NCL-L-CD16 | Novocastra |
| CD163 | clone MRQ-26 # 05973929001 | Ventana. |
| CD1A | clone EP3622 # 06419160001 | Ventana. |
| CD2 | clone MRQ-11 # 05463467001 | Ventana. |
| CD20 | clone L26 # 05267099001 | Ventana. |
| CD23 | clone SP23 # 0547925800 | Ventana. |
| CD25 | clone 4C9 # 05973899001 | Ventana. |
| CD27 | clone 137B4 # NCL-CD27 | Novocastra. |
| CD31 | clone JC70 # 05463475001 | Ventana. |
| CD33 | clone PWS44 # NCL-L-CD33 | Novocastra. |
| CD38 | clone SP149 # 06648550001 | Ventana. |
| CD43 | clone L60 # 05266980001 | Ventana. |
| CD45 | clone RP2/18 # 05266912001 | Ventana. |
| CD56 | clone MRQ-42 # 06433359001 | Ventana. |
| CD61 | clone 2f2 # 05269091001 | Ventana. |
| CD68PG | clone PG-M1 # M0876. Dako | Agilent. |
| CD7 | clone SP94 # 06537847001 | Ventana. |
| CRP | clone Y284 # ab32412 | Abcam. |
| CT1 | polyclonal # ab34710 | Abcam. |
| CT4 | clone CIV 22 # 05267439001 | Ventana. |
| CYCLD1 | clone SP4-R # 05862949001 | Ventana. |
| ECAD | clone 36 # 05905290001 | Ventana. |
| EMA | clone E29 # 05878900001 | Ventana. |
| FVIII | polyclonal # 05267528001 | Ventana. |
| FXIIIA | clone EP3372 # 05922470001 | Ventana. |
| GPA | clone GA-R2 # 05269172001 | Ventana. |
| H2AX | clone Ser139 # 2577 | Cell Signaling. |
| HMGA2 | clone D1A7 # 8179 | Cell Signaling. |
| HSP70 | clone 2A4 # ab5442 | Abcam. |
| IGM | polyclonal # 05267641001 | Ventana. |
| IGG | polyclonal # 05267633001 | Ventana. |
| M-30 | clone M30 # 10700 | Peviva. |
| MPO | polyclonal # 05267692001 | Ventana. |
| MUC1 | clone Ma695 # CM 319 B | Biocare Medical. |
| MUC5AC | clone MRQ-19 # 05463564001 | Ventana. |
| NPM | clone 376 # M7305 | Dako, Agilent. |
| P-SPA | clone RM334 # 31-1221-00 | RevMab Biosciences. |
| TNC | clone T20 # AM20672PU-N | OriGene. |
| PERFORIN | clone MRQ-23 # 05463637001 | Ventana. |
| PDL1 | clone SP263 # 07494190001 | Ventana. |
| PD1 | clone NAT105 # 315M-96 | Cell Marque. |
| FDC | clone CAN.42 # M7157 | Dako, Agilent |
| GRB | clone 11F1 # NCL-L-GRAN-B | Novocastra |
| ACTINM | clone HHF35 # M0635 | Dako, Agilent |
| CK18 | clone DC-10 # M7010 | Dako, Agilent |
| CD146 | clone UMAB154 # UM800051 | Origene |
| CD15 | clone Carb-3 # M3631 | Dako, Agilent |
| CK19 | clone A53-B/A2.26 # 319M-16 | Cell Marque |
| CD10 | clone 56C6 # NCL-L-CD10-270 | Novocastra |
| CD34 | clone QBEnd/10 # M7165 | Dako, Agilent |
| CD30 | clone Ber-H2 # 130M-96 | Cell Marque |
| CD20 | clone L26 # M0755 | Dako, Agilent |
| CD31 | clone JC70 # 131M-96 | Cell Marque |
| ACTINSM | clone 1A4 # M0851 | Dako, Agilent |
| CD3 | clone LN10 # NCL-L-CD3-565 | Novocastra |
| CD4 | clone 4B12 # NCL-L-CD4-368 | Novocastra |
| CD8 | clone C8/144B # M7103 | Dako, Agilent |
| CD56 | clone CD564 # NCL-L-CD56-504 | Novocastra |
| CD68PG | clone PG-M1 # M0876 | Dako, Agilent |
| CK5 | clone XM26 # NCL-L-CK5 | Novocastra |
| CK17 | clone SP95 # ab183330 | Abcam |
| CEAM | clone II-7 # M7072 | Dako, Agilent |
| Podoplanin | clone D2-40 # M3619 | Dako, Agilent |
| KI67 | clone MIB-1 # M7240 | Dako, Agilent |
| Vimentin | clone V9 # M0725 | Dako, Agilent |
| HLA-DR | clone LN-3 # NCL-LN3 | Novocastra |
| P16 | clone G175-405 # 550834 | BD Pharmingen |
| P63 | clone DAK-p63 # M7317 | Dako, Agilent |
| PDGFRA | polyclonal # sc-338 | Santa Cruz |
| PHH3 | polyclonal # 06-570 | Merck |
| PML | clone PG-M3 # sc-966 | Santa Cruz |
| TBM | clone 1009 # M0617 | Dako, Agilent |
| TRAP | clone 26E5 # NCL-L-TRAP | Novocastra |
| TTF1 | clone SPT24 # NCL-L-TTF-1 | Novocastra |
| NAPSINA | clone MRQ-60 # 352M-96 | Cell Marque |
| CHROMA | clone LK2H10 # 238M-96 | Cell Marque |
| P40 | polyclonal # ACI 3030B | Biocare Medical |

## References

Ackermann, M., Verleden, S. E., Kuehnel, M., Haverich, A., Welte, T., Laenger, F., Vanstapel, A., Werlein, C. M. D., Stark, H., Tzankov, A., Li, W. W., Li, V. W., Mentzer, S. J., & Jonigk, D. (2020). Pulmonary vascular endothelialitis, thrombosis, and angiogenesis in Covid-19. New England Journal of Medicine, 383(2), 120–128. https://doi.org/10.1056/NEJMoa2015432

Adachi, T., Chong, J.-M., Nakajima, N., Sano, M., Yamazaki, J., Miyamoto, I., Nishioka, H., Akita, H., Sato, Y., Kataoka, M., Katano, H., Tobiume, M., Sekizuka, T., Itokawa, K., Kuroda, M., & Suzuki, T. (2020). Clinicopathologic and Immunohistochemical Findings from Autopsy of Patient with COVID-19, Japan. Emerging Infectious Diseases, 26(9). https://doi.org/10.3201/eid2609.201353

Arvin, A. M., Fink, K., Schmid, M. A., Cathcart, A., Spreafico, R., Havenar-Daughton, C., Lanzavecchia, A., Corti, D., & Virgin, H. W. (2020). A perspective on potential antibody-dependent enhancement of SARS-CoV-2. Nature, 584(August), 353–363. https://doi.org/10.1038/s41586-020-2538-8

Barnes, B. J., Adrover, J. M., Baxter-Stoltzfus, A., Borczuk, A., Cools-Lartigue, J., Crawford, J. M., Daßler-Plenker, J., Guerci, P., Huynh, C., Knight, J. S., Loda, M., Looney, M. R., McAllister, F., Rayes, R., Renaud, S., Rousseau, S., Salvatore, S., Schwartz, R. E., Spicer, J. D., … Egeblad, M. (2020). Targeting potential drivers of COVID-19: Neutrophil extracellular traps. Journal of Experimental Medicine, 217(6), 1–7. https://doi.org/10.1084/jem.20200652

Barton, L. M., Duval, E. J., Stroberg, E., Ghosh, S., & Mukhopadhyay, S. (2020). COVID-19 Autopsies, Oklahoma, USA. American Journal of Clinical Pathology, 153(6), 725–733. https://doi.org/10.1093/ajcp/aqaa062

Becker, R. C. (2020). COVID-19-associated vasculitis and vasculopathy. Journal of Thrombosis and Thrombolysis, 50(3), 499–511. https://doi.org/10.1007/s11239-020-02230-4

Bradley, B. T., Maioli, H., Johnston, R., Chaudhry, I., Fink, S. L., Xu, H., Najafian, B., Deutsch, G., Lacy, J. M., Williams, T., Yarid, N., & Marshall, D. A. (2020). Histopathology and ultrastructural findings of fatal COVID-19 infections in Washington State: a case series. The Lancet, 396(10247), 320–332. https://doi.org/10.1016/s0140-6736(20)31305-2

Calabrese, F., Pezzuto, F., Fortarezza, F., Hofman, P., Kern, I., Panizo, A., von der Thüsen, J., Timofeev, S., Gorkiewicz, G., & Lunardi, F. (2020). Pulmonary pathology and COVID-19: lessons from autopsy. The experience of European Pulmonary Pathologists. Virchows Archiv, December 2019, 359–372. https://doi.org/10.1007/s00428-020-02886-6

Chan, C. M., Tsoi, H., Chan, W. M., Zhai, S., Wong, C. O., Yao, X., Chan, W. Y., Tsui, S. K. W., & Chan, H. Y. E. (2009). The ion channel activity of the SARS-coronavirus 3a protein is linked to its pro-apoptotic function. International Journal of Biochemistry and Cell Biology, 41(11), 2232–2239. https://doi.org/10.1016/j.biocel.2009.04.019

Chan, K. H., Peiris, J. S. M., Lam, S. Y., Poon, L. L. M., Yuen, K. Y., & Seto, W. H. (2011). The effects of temperature and relative humidity on the viability of the SARS coronavirus. Advances in Virology, 2011. https://doi.org/10.1155/2011/734690

Chen, L., Zhong, Y., Sheng, G., & Zou, L. (2020). Koch’s time for COVID-19 and the other major epidemics of the 21st century. 1–9. https://doi.org/10.31219/osf.io/wsqcf

Doremalen, Bushmaker, M. (2020). Aerosol and Surface Stability of SARS-CoV-2 as Compared with SARS-CoV-1 To. The New England Journal of Medicine, March 27.

Edler, C., Schröder, A. S., Aepfelbacher, M., Fitzek, A., Heinemann, A., Heinrich, F., Klein, A., Langenwalder, F., Lütgehetmann, M., Meißner, K., Püschel, K., Schädler, J., Steurer, S., Mushumba, H., & Sperhake, J. P. (2020). Dying with SARS-CoV-2 infection—an autopsy study of the first consecutive 80 cases in Hamburg, Germany. International Journal of Legal Medicine, 134(4), 1275–1284. https://doi.org/10.1007/s00414-020-02317-w

Eroshenko, N., Gill, T., Keaveney, M. K., Church, G. M., Trevejo, J. M., & Rajaniemi, H. (2020). Implications of antibody-dependent enhancement of infection for SARS-CoV-2 countermeasures. Nature Biotechnology, 38(7), 789–791. https://doi.org/10.1038/s41587-020-0577-1

Hanley, B., Naresh, K. N., Roufosse, C., Nicholson, A. G., Weir, J., Cooke, G. S., Thursz, M., Manousou, P., Corbett, R., Goldin, R., Al-Sarraj, S., Abdolrasouli, A., Swann, O. C., Baillon, L., Penn, R., Barclay, W. S., Viola, P., & Osborn, M. (2020). Histopathological findings and viral tropism in UK patients with severe fatal COVID-19: a post-mortem study. In The Lancet. Microbe. https://doi.org/10.1016/S2666-5247(20)30115-4

He, L., Mäe, M. A., Sun, Y., Muhl, L., Nahar, K., Liébanas, E. V., Fagerlund, M. J., Oldner, A., Liu, J., Genové, G., Pietilä, R., Zhang, L., Xie, Y., Leptidis, S., Mocci, G., Stritt, S., Osman, A., Anisimov, A., Hemanthakumar, K. A., … Betsholtz, C. (2020). Pericyte-specific vascular expression of SARS-CoV-2 receptor ACE2 – implications for microvascular inflammation and hypercoagulopathy in COVID-19 patients. BioRxiv, 2020.05.11.088500. https://doi.org/10.1101/2020.05.11.088500

Huang, C., Wang, Y., Li, X., Ren, L., Zhao, J., Hu, Y., Zhang, L., Fan, G., Xu, J., Gu, X., Cheng, Z., Yu, T., Xia, J., Wei, Y., Wu, W., Xie, X., Yin, W., Li, H., Liu, M., … Cao, B. (2020). Clinical features of patients infected with 2019 novel coronavirus in Wuhan, China. The Lancet, 6736(20), 1–10. https://doi.org/10.1016/s0140-6736(20)30183-5

Hui, K. P. Y., Cheung, M. C., Perera, R. A. P. M., Ng, K. C., Bui, C. H. T., Ho, J. C. W., Ng, M. M. T., Kuok, D. I. T., Shih, K. C., Tsao, S. W., Poon, L. L. M., Peiris, M., Nicholls, J. M., & Chan, M. C. W. (2020). Tropism, replication competence, and innate immune responses of the coronavirus SARS-CoV-2 in human respiratory tract and conjunctiva: an analysis in ex-vivo and in-vitro cultures. The Lancet Respiratory Medicine, 8(7), 687–695. https://doi.org/10.1016/S2213-2600(20)30193-4

Kipar, A., & Meli, M. L. (2014). Feline Infectious Peritonitis: Still an Enigma? Veterinary Pathology, 51(2), 505–526. https://doi.org/10.1177/0300985814522077

Kowalik, M. M., Trzonkowski, P., Łasińska-Kowara, M., Mital, A., Smiatacz, T., & Jaguszewski, M. (2020). COVID-19 — Toward a comprehensive understanding of the disease. Cardiology Journal, 27(2), 99–114. https://doi.org/10.5603/CJ.a2020.0065

Kox, M., Waalders, N. J. B., Kooistra, E. J., Gerretsen, J., & Pickkers, P. (2020). Cytokine Levels in Critically Ill Patients With COVID-19 and Other Conditions. JAMA. https://doi.org/10.1001/jama.2020.17052

Liu, X., Zhu, A., He, J., Chen, Z., Liu, L., Xu, Y., Ye, F., Feng, H., Luo, L., Cai, B., Mai, Y., Lin, L., Zhang, Z., Chen, S., Shi, J., Wen, L., Wei, Y., Zhuo, J., Zhao, Y.,… Chen, J. (2020). Single-Cell Analysis Reveals Macrophage-Driven T Cell Dysfunction in Severe COVID-19 Patients. MedRxiv, 2, 2020.05.23.20100024-2020.05.23.20100024. https://doi.org/10.1101/2020.05.23.20100024

Loomba, R. S., Villarreal, E. G., & Flores, S. (2020). COVID-19 and Hyperinflammatory Syndrome in Children: Kawasaki Disease with Macrophage Activation Syndrome in Disguise? Cureus, 12(8), 4–10. https://doi.org/10.7759/cureus.9515

Luo, F., Liao, F. L., Wang, H., Tang, H. Bin, Yang, Z. Q., & Hou, W. (2018). Evaluation of Antibody-Dependent Enhancement of SARS-CoV Infection in Rhesus Macaques Immunized with an Inactivated SARS-CoV Vaccine. Virologica Sinica, 33(2), 201–204. https://doi.org/10.1007/s12250-018-0009-2

Martines, R. B., Ritter, J. M., Matkovic, E., Gary, J., Bollweg, B. C., Bullock, H., Goldsmith, C. S., Silva-Flannery, L., Seixas, J. N., Reagan-Steiner, S., Uyeki, T., Denison, A., Bhatnagar, J., Shieh, W.-J., & Zaki, S. R. (2020). Pathology and Pathogenesis of SARS-CoV-2 Associated with Fatal Coronavirus Disease, United States. Emerging Infectious Diseases, 26(9), 2005–2015. https://doi.org/10.3201/eid2609.202095

McElroy, A. K., Shrivastava-Ranjan, P., Harmon, J. R., Martines, R. B., Silva-Flannery, L., Flietstra, T. D., Kraft, C. S., Mehta, A. K., Lyon, G. M., Varkey, J. B., Ribner, B. S., Nichol, S. T., Zaki, S. R., & Spiropoulou, C. F. (2019). Macrophage activation marker soluble cd163 associated with fatal and severe ebola virus disease in humans. Emerging Infectious Diseases, 25(2), 290–298. https://doi.org/10.3201/eid2502.181326

Molina, D. K., & DiMaio, V. J. M. (2015). Normal Organ Weights in Women: Part II-The Brain, Lungs, Liver, Spleen, and Kidneys. The American Journal of Forensic Medicine and Pathology, 36(3), 182–187. https://doi.org/10.1097/paf.0000000000000175

Mucha, S. R., Dugar, S., McCrae, K., Joseph, D. E., Bartholomew, J., Sacha, G., & Militello, M. (2020). Coagulopathy in COVID-19: Posted april 24, 2020. Cleveland Clinic Journal of Medicine, 87(5), 1–6. https://doi.org/10.3949/CCJM.87A.CCC024

Peeples, L. (2020). Avoiding pitfalls in the pursuit of a COVID-19 vaccine. Proceedings of the National Academy of Sciences of the United States of America, 117(15), 8218–8221. https://doi.org/10.1073/pnas.2005456117

Ren, Y., Shu, T., Wu, D., Mu, J., Wang, C., Huang, M., Han, Y., Zhang, X. Y., Zhou, W., Qiu, Y., & Zhou, X. (2020). The ORF3a protein of SARS-CoV-2 induces apoptosis in cells. Cellular and Molecular Immunology, 17(8), 881–883. https://doi.org/10.1038/s41423-020-0485-9

Salah, H. M., Naser, J. A., Calcaterra, G., Bassareo, P. P., & Mehta, J. L. (2020). The Effect of Anticoagulation Use on Mortality in COVID-19 Infection. In American Journal of Cardiology. https://doi.org/10.1016/j.amjcard.2020.08.005

Salerno, M., Sessa, F., Piscopo, A., Montana, A., Torrisi, M., Patanè, F., Murabito, P., Li Volti, G., & Pomara, C. (2020). No Autopsies on COVID-19 Deaths: A Missed Opportunity and the Lockdown of Science. Journal of Clinical Medicine, 9(5), 1472. https://doi.org/10.3390/jcm9051472

Schulte-Schrepping, J., Reusch, N., Paclik, D., Baßler, K., Schlickeiser, S., Zhang, B., Krämer, B., Krammer, T., Brumhard, S., Bonaguro, L., De Domenico, E., Wendisch, D., Grasshoff, M., Kapellos, T. S., Beckstette, M., Pecht, T., Saglam, A., Dietrich, O., Mei, H. E., … Ziebuhr, J. (2020). Severe COVID-19 Is Marked by a Dysregulated Myeloid Cell Compartment. Cell. https://doi.org/10.1016/j.cell.2020.08.001

Srikiatkhachorn, A., & Yoon, I. K. (2016). Immune correlates for dengue vaccine development. Expert Review of Vaccines, 15(4), 455–465. https://doi.org/10.1586/14760584.2016.1116949

Suess, C., & Hausmann, R. (2020). Gross and histopathological pulmonary findings in a COVID-19 associated death during self-isolation. International Journal of Legal Medicine, 134(4), 1285–1290. https://doi.org/10.1007/s00414-020-02319-8

Takano, T., Kawakami, C., Yamada, S., Satoh, R., & Hohdatsu, T. (2008). Antibody-dependent enhancement occurs upon re-infection with the identical serotype virus in feline infectious peritonitis virus infection. Journal of Veterinary Medical Science, 70(12), 1315–1321. https://doi.org/10.1292/jvms.70.1315

Varga, Z., Flammer, A. J., Steiger, P., Haberecker, M., Andermatt, R., Zinkernagel, A. S., Mehra, M. R., Schuepbach, R. A., Ruschitzka, F., & Moch, H. (2020). Endothelial cell infection and endotheliitis in COVID-19. The Lancet, 395(10234), 1417–1418. https://doi.org/10.1016/S0140-6736(20)30937-5

Vasquez-Bonilla, W. O., Orozco, R., Argueta, V., Sierra, M., Zambrano, L. I., Muñoz-Lara, F., López-Molina, D. S., Arteaga-Livias, K., Grimes, Z., Bryce, C., Paniz-Mondolfi, A., & Rodríguez-Morales, A. J. (2020). A review of the main histopathological findings in coronavirus disease 2019. Human Pathology. https://doi.org/10.1016/j.humpath.2020.07.023

Wichmann, D., Sperhake, J.-P., Lütgehetmann, M., Steurer, S., Edler, C., Heinemann, A., Heinrich, F., Mushumba, H., Kniep, I., Schröder, A. S., Burdelski, C., de Heer, G., Nierhaus, A., Frings, D., Pfefferle, S., Becker, H., Bredereke-Wiedling, H., de Weerth, A., Paschen, H.-R., … Kluge, S. (2020a). Autopsy Findings and Venous Thromboembolism in Patients With COVID-19. Annals of Internal Medicine, 25(4). https://doi.org/10.7326/m20-2003

Wichmann, D., Sperhake, J.-P., Lütgehetmann, M., Steurer, S., Edler, C., Heinemann, A., Heinrich, F., Mushumba, H., Kniep, I., Schröder, A. S., Burdelski, C., de Heer, G., Nierhaus, A., Frings, D., Pfefferle, S., Becker, H., Bredereke-Wiedling, H., de Weerth, A., Paschen, H.-R., … Kluge, S. (2020b). Autopsy Findings and Venous Thromboembolism in Patients With COVID-19. Annals of Internal Medicine, 25(4). https://doi.org/10.7326/m20-2003

Zhou, B., Zhao, W., Feng, R., Zhang, X., Li, X., Zhou, Y., Peng, L., Li, Y., Zhang, J., Luo, J., Li, L., Wu, J., Yang, C., Wang, M., Zhao, Y., Wang, K., Yu, H., Peng, Q., & Jiang, N. (2020). The pathological autopsy of coronavirus disease 2019 (COVID-2019) in China: a review. Pathogens and Disease, 78(3), 1–9. https://doi.org/10.1093/femspd/ftaa026

Zhou Baoyong, Zhao Wei, Feng, Ruixi, Zhang Xiaohui, Li Xuemei, Zhou Yang, Peng Li, Li Yixin, J., Zhang, Luo Jing, L. L., & WuJingxian, Yang Changhong, Wang Meijiao, Zhao Yong, Wang Kejian, Yu Huarong, Peng Qiling, J. N. (2020). The pathologic autopsy of coronavirus disease 2019 (COVID-2019) in China : a review. Pathogens and Disease, 78(3). https://doi.org/10.1093/femspd/ftaa026

